# Bingeing in rats: Persistence of high intakes of palatable solutions induced by 1-in-4 days intermittent access

**DOI:** 10.1101/539742

**Authors:** Simone Rehn, Robert A. Boakes

**Affiliations:** School of Psychology, The University of Sydney

**Author notes:** Correspondence: Professor R. A. Boakes, School of Psychology, University of Sydney, NSW 2006, Australia.

**Keywords:** Bingeing, sucrose, saccharin, maltodextrin, rats

## Abstract

When animals are given access to a palatable food or drink on some days but not on others, the amount they consume can far exceed the daily amounts consumed by controls given daily access. In a previous study such bingeing was found when rats were given 4% sucrose solution; it also found that, following 1-in-4-days access for many weeks, intakes remained persistently higher than that of controls even when the conditions were changed to 1-in-2-days access for both groups. One aim of the three experiments reported here was to test whether such persistent bingeing could be found for other solutions. This was confirmed in rats for a saccharin solution and a highly palatable saccharin-plus-glucose solution. However, when a maltodextrin solution was used, initial increased intakes produced by the 1-in-4-days schedule were not maintained when this was changed to a 1-in-2-days schedule. These results suggested that the hedonic value of a solution is more important than its caloric content in determining whether it will support persistent bingeing. A second aim was to test for evidence that the 1-in-4-days procedure induced an addiction to the target solution. No such evidence was found using multiple measures including instrumental responding and anxiety-like behavior on the elevated plus-maze for craving and withdrawal respectively.

## 1. Introduction

Binge eating can be defined as the consumption of a large amount of food in a short period of time (APA, 2013). In humans it can lead to detrimental consequences to individuals’ physical and psychological well-being. For example, binge eating is associated with a loss of control, intense guilt, and excessive weight gain over the long-term, especially when binges occur without compensation at other times for an increased caloric intake (i.e. binge-eating disorder) (APA, 2013). The types of food consumed in binges tend to be those high in sugar and fat (Yanovski et al., 1992), the overconsumption of which have been linked to obesity and cognitive impairments in human studies (Francis & Stevenson, 2011; Kanoski & Davidson, 2011; Nyaradi et al., 2014).

Although many binge-eating individuals acknowledge the associated health problems (Colles, Dixon, & O’Brien, 2008) and experience distress over their binge eating (APA, 2013), they continue engaging in compulsive eating behaviors. This loss of control highlights the difficulty of treatment upon the onset of problematic binge eating and emphasizes the need to understand factors underlying binge-eating development. Several animal models have consequently been established to explore the development of binge-like consumption of high-sugar and/or high-fat food and drinks. Some of these models provide access to highly palatable foods or drinks for a limited period each day (e.g. Avena, Rada, & Hoebel, 2008). Arguably, these fail to model typical human bingeing behavior and are subject to confounding by circadian entrainment (e.g. Eikelboom & Hewitt, 2016). Others provide such access on only certain days (e.g. Corwin & Wojnicki, 2006), a pattern that resembles human bingeing (Kales, 1990).

A common finding in such animal studies is that intakes during periods of intermittent access are far greater than the average intakes during similar periods by animals with unrestricted access to the same highly palatable foods or drinks (e.g. Avena, Rada, et al., 2008; Corwin & Wojnicki, 2006). Such a result was also reported by Eikelboom and Hewitt (2016). In their series of experiments intermittent access to a sucrose solution produced long-term increases in consumption that resembled bingeing. What was remarkable about their study was that under some conditions these elevated intakes persisted even when the conditions that produced them were terminated. In their first experiment, rats were given either continuous, second-, third-, or fourth-day 23.5-h access to a 4% sucrose solution for 49 days (Phase 1) before all were switched to alternate-day access for an additional 24 days (Phase 2). The most striking results were obtained from the intermittent group given sucrose solution every fourth day; these rats came to consume up to three times the amount of sucrose solution (~300 g) in 23.5 h relative to rats with continuous access (~100 g) in Phase 1. Critically, when shifted to identical alternate-day access conditions in Phase 2, the fourth-day-access group maintained much higher sucrose intakes relative to the continuous group for the remainder of the experiment. Since the largest intake difference was found between these two groups, they will hereafter be referred to here as the Binge and Unrestricted groups of the Eikelboom protocol.

When compared with previous binge models (e.g. Avena, Rada, et al., 2008; Corwin & Wojnicki, 2006), Eikelboom and Hewitt (2016) appears to be the only study to demonstrate *persistence* of elevated intakes induced by intermittent-access conditions. Additionally, this protocol specifies that the deprivation condition of the binge (fourth-day access; Binge) and non-binge (continuous access; Unrestricted) animals is identical with regards to sucrose access during Phase 2, yet the binge animals repeatedly consume much larger amounts in the same period of time, thus satisfying the operationalization of a binge (Corwin & Buda-Levin, 2004). This intake difference also mimics the criterion of ‘objectively larger amounts’ consumed in human binge episodes (APA, 2013). Finally, rats were fed *ad libitum* in this protocol, which allows a distinction between binge-like and homeostatic consumption.

The Eikelboom protocol also eliminates circadian entrainment effects by providing 24-h access to a sucrose solution on the day that it is available. This extended access is vital for producing the absolute increases in daily sucrose intake seen in the binge group. Procedures that maintain circadian regularity usually fail to produce differences in total daily intake between binge and non-binge groups (Avena, Rada, et al., 2008). In the case of fat bingeing rats given access schedules that maintain circadian regularity (i.e. 2-h daily fat access) exhibit much smaller elevations in intake compared to rats that do not (i.e. 2-h fat access on three days a week). Furthermore, the elimination of circadian entrainment effects is essential to avoid artefactual patterns of behavior. The alignment of circadian rhythm with periodic time cues, such as light/dark cycles or expected feeding times, produces daily rhythms of food-anticipatory activity before expected meal times (Mistlberger, 1993). This activity can manifest as increased wheel running (Bolles & Stokes, 1965) and lever pressing (Boulos, Rosenwasser, & Terman, 1980). However, food-anticipatory activity does not occur in rats fed *ad libitum* (Landry, Yamakawa, Webb, Mear, & Mistlberger, 2007) nor with day-long sucrose access that does not align with circadian rhythm.

By avoiding circadian-entrained food-anticipatory behavior, this protocol also enables a clearer examination of the overlap between bingeing and addiction (see Corwin & Babbs, 2012 for review). Hoebel’s influential model of ‘sugar addiction’ has proposed that intermittent, excessive intake of sugar produces ‘withdrawal’ (Colantuoni et al., 2002) and ‘craving’ (Avena, Long, & Hoebel, 2005). However, this model pits food-entrained rhythms against light-entrained rhythms; rats are food-deprived daily for 12 h and then given 12-h access to sugar (25% glucose or 10% sucrose) and chow 4 h into the dark cycle of their circadian rhythm (Avena, Rada, et al., 2008). Under these competing conditions, access to sugar can engage food-entrained rhythms and result in elevated activity during expected sugar-access times (Bolles & Stokes, 1965; Pecoraro, Gomez, Laugero, & Dallman, 2002). It has been argued that findings of increased lever-pressing in sugar-bingeing rats relative to controls after abstinence is an indication of ‘craving’ (Avena et al., 2005; Avena, Rada, et al., 2008). It is unclear, however, whether the control group in this study were under similar food deprivation conditions as the sugar-bingeing rats. Given that food-anticipatory activity would not occur in controls fed *ad libitum* (Landry et al., 2007), increased lever-pressing in the sugar-bingeing rats may instead be attributed to food-anticipatory behavior. Similarly, sugar-bingeing rats may have exhibited greater anxiety-like behavior on an elevated plus-maze (EPM) than an *ad libitum* chow group – which has been taken to indicate ‘withdrawal’ – because they were denied access to chow and sugar during expected feeding times (Colantuoni et al., 2002). This study included a cyclic glucose control group which would have also displayed food-anticipatory behavior, but whether the sugar-bingeing group differed from these controls was not reported (Colantuoni et al., 2002).

Given the advantages of the Eikelboom protocol in promoting binge-like sucrose consumption in rats, the present study had two aims: First, to test whether the persistence of binge-like consumption of sucrose would generalize to similarly attractive solutions and, second, to test whether persistent bingeing would be accompanied by addiction-like behavior. To our knowledge persistent elevation in the consumption of solutions other than 4% sucrose has not been examined. Maltodextrin is an example of a non-sweet polysaccharide that has similar metabolic effects to sucrose in rats (Kendig, Lin, Beilharz, Rooney, & Boakes, 2014; Nissenbaum & Sclafani, 1987). Rats readily consume maltodextrin and are able to discriminate its taste from sucrose, preferring maltodextrin over sucrose at low concentrations (Sclafani, 1987). Furthermore, adding saccharin to either sucrose, glucose or maltodextrin solutions produces a polydipsic effect; rats find such mixed solutions highly palatable and prefer them to the solutions presented alone (Sclafani, Einberg, & Nissenbaum, 1987). Therefore, to meet the first aim the current study examined whether persistent binge-like sucrose consumption under the Eikelboom protocol could be replicated under somewhat different conditions (Experiment 1) and whether it would generalize to similarly attractive target solutions. In Experiment 2 this was sweet, yet non-caloric, saccharin and caloric, yet non-sweet, maltodextrin solutions. In Experiment 3 a highly-palatable mixture of saccharin and glucose was compared to saccharin solution alone.

The persistence of elevated intakes in Eikelboom and Hewitt (2016)’s study resemble tolerance in addiction, which highlight its relevance to exploring the overlap between bingeing and addiction (see Corwin & Babbs, 2012 for review). Therefore, in addition to examining whether persistent elevations would generalize to other palatable solutions, the current study also explored whether addiction-like behavior such as ‘withdrawal’ and ‘craving’ would arise as a result of binge-like consumption of these solutions. To this end, several behavioral measures were employed. These consisted of: 1) lever-press responding under a variable-ratio (VR) schedule; 2) a flavor preference test; 3) a preference test between the target solution and an equally attractive solution; and 4) withdrawal-induced anxiety-like behavior on the elevated plus maze (EPM).

The first three tests were measures of ‘craving’, defined as ‘the incentive motivation to self-administer an abused substance or respond for its associated cues’ (Markou et al., 1993). Variable Ratio (VR) reinforcement schedules have been shown to be sensitive to changes in self-administration behavior with sucrose reinforcers (Petry & Heyman, 1995). Thus, VR schedules were used to test whether motivation to respond for a target solution reinforcer differed between Binge and Unrestricted groups.

In the flavor preference test, a novel flavor (i.e. almond) is initially paired with the target solution (e.g. sucrose) and preference for this flavor over a flavorless solution is taken as a measure of incentive salience. Flavor preference learning is based on the ability of sucrose and other highly palatable substances to impart conditioned incentive value onto previously neutral stimuli, akin to drug-related cues which come to elicit drug-taking behavior (Markou et al., 1993). Use of this measure was based on a parallel with salt craving: Inducing a sodium deficiency in rats increases their preference for a flavor previously paired with salt (Fudim, 1978). This flavor preference test was previously used in a study that found sucrose-bingeing rats in an adapted Hoebel model to exhibit increased preference for a sucrose-associated almond flavor (Wu & Boakes, in preparation).

An additional preference test between the target solution (e.g. sucrose) and an equally attractive solution (e.g. maltodextrin) was employed as a further measure of craving. This measures the unconditioned incentive value, or the reinforcing properties (i.e. hedonic value) of the target solution itself (Markou et al., 1993). Given that an increase in hedonic set point has been implicated in addiction (Ahmed & Koob, 1998), this preference test aimed to examine whether Binge groups would show a greater preference for the target solution relative to an isohedonic solution, reflecting its increased hedonic value, after engaging in persistent binge-like consumption of the target solution.

In accordance with existing animal models of drug addiction (Schulteis, Yackey, Risbrough, & Koob, 1998; Walf & Frye, 2007), withdrawal in the present study was operationalized as anxiety-like behavior, as measured on the EPM, following a period when the target solution was no longer available. A smaller proportion of time spent on the open arms can indicate greater anxiety in the rat (Walf & Frye, 2007).

## 1. Experiment 1: Sucrose bingeing, withdrawal, and craving

Experiment 1 aimed both to confirm the previous finding that giving rats every-fourth-day access to 4% sucrose solution (Binge group) can produce long-lasting elevations in sucrose intake even when switched to alternate-day access (Eikelboom & Hewitt, 2016) and to determine whether such binge-like sucrose consumption can produce withdrawal and craving. Eikelboom and Hewitt (2016) reported the persistence effect in their Experiment 1 following a 49-day Phase 1 but not in their Experiment 2, where Phase 1 lasted only 10 days. An intermediate length of Phase 1 (28 days) was used in the present experiment. Two control groups were included: An Unrestricted group given access to 4% sucrose daily in Phase 1 and a Chow group that received only chow and water throughout. The latter served to clarify whether prolonged sucrose exposure in the Unrestricted group – independent of bingeing – would affect weight, chow intake, and performance on behavioral measures of withdrawal and craving.

Bingeing was operationalized as: 1) A greater escalation of intake across time in Phase 1, which is an indicator of bingeing in both addiction and binge models (Corwin & Babbs, 2012) and; 2) greater sucrose consumption in a 24-h period by Binge rats than by Unrestricted rats under identical access conditions in Phase 2. Following the proposed relationship between bingeing and addiction (Avena, Rada, et al., 2008; Corwin & Babbs, 2012), the Binge group was predicted to demonstrate greater withdrawal and craving than the Unrestricted group.

### 2.1. Methods

#### 2.1.1. Animals

Thirty male Sprague-Dawley rats were purchased from the Animal Resource Centre (ARC), Perth. They were six weeks old, with an average weight of 221 g (range 200 – 254 g), on arrival, when they were initially group-housed (*n* = 5/cage) in open-topped cages (59 × 36 × 19cm). The temperature- and humidity-controlled colony room was maintained on a reversed 12-h light/dark cycle (lights off at 0800 hrs). On completion of the Pre-diet tests rats were transferred to single housing in open-topped shoebox cages (47 × 32 × 14 cm) to allow monitoring of individual chow and fluid intakes throughout the rest of the experiment. Body weight, chow intake, and water intake were measured every four days throughout the experiment. Target solution intakes were measured before and after the Binge rats’ access day in Phase 1 and daily in Phase 2. Cage bedding was changed once or twice a week. Tap water (Sydney Water) and chow (Specialty Feeds^®^, 14.2 kJ/g, Glen Forrest, WA) were available *ad libitum* throughout unless otherwise noted. All procedures were approved by the University of Sydney Animal Ethics Committee.

#### 2.1.2. Solutions

All target solutions were mixed based on a weight/volume (w/v) basis using tap water (Sydney Water). Sucrose solutions were mixed using commercially-available pure cane sugar (17 kJ/g). Maltodextrin solutions were mixed using maltodextrin (16kJ/g, Myopure Maltodextrin DE17; www.myopure.com.au). Almond-flavored solutions were mixed on a volume/volume (v/v) basis using almond essence (Queen).

#### 2.1.3 Apparatus

Ten operant chambers (MED Associates, East Fairfield, VT) were contained within sound-attenuated and ventilated cubicles. Each chamber contained two levers located on either side of the magazine, and the lever to the left of the magazine was active. Each active lever press produced 10-s access to 0.1 mL of 4% sucrose solution, delivered via a retractable dipper. Dipper presentations were accompanied by a 1-s tone and the chamber light turning off, indicating reinforcer availability. The magazine recesses contained infrared sensors that detected nose pokes. LabVIEW software (National Instruments, Austin, TX) controlled reinforcement schedules in these chambers.

For preference training and tests, ten acrylic cages (36 × 20 × 18 cm) fitted with lids and paper-pellet bedding were used as individual drinking chambers. 100-mL plastic bottles with ball-bearing stainless steel spouts contained drinking solutions and were inserted into the cages.

The elevated plus-maze (EPM) was composed of four arms (11 × 45 cm) intersecting at a central open square (10 × 10 cm) and elevated 80 cm above the floor. Two opposite arms (closed arms) were enclosed with opaque walls (40 cm high), and the other two arms (open arms) had no walls. During each session, the animals’ behavior was recorded using a video camera mounted at a height of 1.15 m vertically above the center of the EPM.

#### 2.1.4 Procedure

An outline of the procedure for Experiment 1 is summarized in Table 1.

**Table 1.**
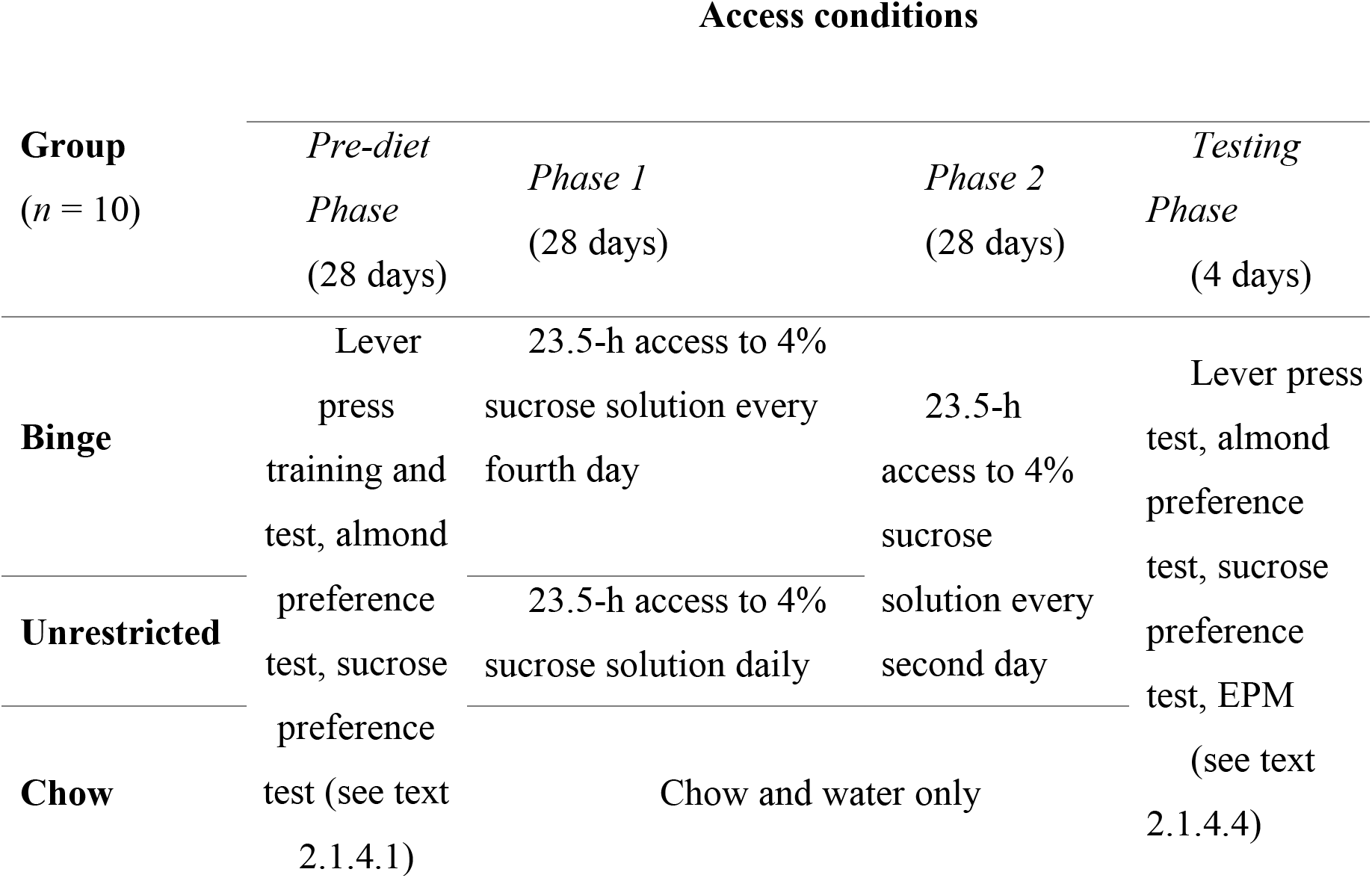
Design of Experiment 1.

##### 2.1.4.1. Pre-diet Phase (Days 1-28)

After five days of acclimatization and handling, the pre-diet phase began with chow restricted to 85 g per group cage per day, given after daily lever-press sessions and with water always available, except for 1 h before sessions.

*Lever press training (Days 1-28)*. A 4% (w/v) sucrose solution reinforcer was used during training and test sessions throughout Experiment 1. Each magazine training and lever-press-training session lasted 30 min. Rats received two magazine training sessions where dipper presentations were on a fixed time (FT-30s) schedule. This resulted in successful magazine-training for all 30 rats, according to the criterion of making at least five magazine entries per session in both sessions.

Each rat was then given lever-press training using continuous reinforcement (FR-1).

Training was considered successful when a rat made at least 25 lever presses within a session. Up to 18 sessions were given, and data from rats failing this criterion were excluded from consequent analyses (*n* = 6). Each rat then received four lever-press training sessions across four consecutive days (one session per day) using the following reinforcement schedules: VI-10s, VI-10s, VR-5, VR-5, where VI indicates a variable-interval and VR a variable-ratio schedule.

*Almond preference training and test (Days 10-12, 21)*. All preference training and test sessions lasted 10 min. Each rat received three training sessions across three consecutive days (one session per day) and one test session. Bottles were weighed before and after each session to calculate consumption to the nearest 0.1g.

For the initial training session, each rat was given a single bottle containing a 4% sucrose + 1% (v/v) almond solution. In the second and third sessions, each rat was given two bottles both containing the same sucrose + almond solution, and the positions of the bottles were exchanged after 5 min to acclimate rats to the two-bottle choice test procedure.

In the two-bottle choice test, a base solution of 1% (w/v) sucrose was used to ensure sufficient fluid consumption in both bottles, such that each rat was given a choice between one bottle containing the base solution (1% sucrose) and another bottle containing the base + almond solution (1% sucrose + 1% almond). The bottle positions were exchanged after 5 min. The initial position of the bottle containing the base + almond solution was counterbalanced between groups. Almond preference was calculated as the consumption of the base + almond solution as a percentage of total fluid consumption (base + almond and base solution) in the two-bottle choice tests.

*Sucrose preference test (Day 25)*. Prior to this test, all rats were given overnight access to 4% maltodextrin in their home cages to reduce a potential neophobic response in the subsequent test. The following day, each rat was given a two-bottle choice test between 4% sucrose and 4% maltodextrin using the procedure described previously for the almond preference test. In the first 5 min of each session, 4% maltodextrin was placed on the right and 4% sucrose was on the left. Sucrose preference was calculated as the consumption of 4% sucrose solution as a percentage of total fluid consumption (4% maltodextrin and 4% sucrose) in the two-bottle choice tests.

##### 2.1.4.2. Phase 1 (Days 29-56)

Rats were allocated to three groups (*n* = 10/group) matched for body weight, almond preference and baseline lever-press responding. Over the 28 days of this phase the Binge group received 23.5-h access to 4% sucrose every fourth day, starting at 0930 hrs and ending at 0900 hrs the next day, while the Unrestricted group received 23.5-h access to 4% sucrose daily. The Chow group were maintained on chow and water and never received sucrose access in the home cages.

##### 2.1.4.3. Phase 2 (Days 57-84)

Binge and Unrestricted groups were switched onto an alternate-day access schedule and given 23.5-h access to 4% sucrose every second day, starting at 0930 hrs and ending at 0900 hrs the next day. The Chow group remained on chow and water only.

Lever-press tests were conducted during Phase 2 on days that rats did not receive access to sucrose (non-sucrose days). On two non-sucrose days at the beginning (Days 58, 60) and end of Phase 2 (Days 82, 84), chow was removed 3 h (at 0900 hrs) before each lever-press test session. At each time point, each rat was tested on a VI-10s schedule for 4% sucrose solution on the first non-sucrose day and on a VR-5 schedule on the next non-sucrose day.

Immediately following the VI-10s lever-press sessions (Days 58, 82) rats were tested for their almond preference using the two-bottle choice test previously described. Immediately following the VR-5 lever-press sessions (Days 60, 84) rats were tested for their preference for sucrose over maltodextrin.

##### 2.1.4.4 Testing Phase (Days 85-88)

On Day 85 five rats from each group (Non-staggered) were sucrose-deprived for 48 h, after which they underwent EPM testing (Day 86). The remaining rats (Staggered) received an extra day of sucrose access on Day 85, and similarly underwent EPM testing after 48-h sucrose-deprivation (Day 88). Each rat was tested on the EPM for 5 min, starting with an initial placement in the center of the EPM facing an open arm. The EPM was wiped down with 50% (v/v) ethanol after each rat.

All 30 EPM video recordings were scored by a non-blind experimenter, and the time each rat spent in the open arms, closed arms and central square was recorded. Behavior on the EPM was calculated as time spent on the open arms as a percentage of the total time spent on both arms. To establish inter-rater reliability, 15 of these recordings were rated by a blind scorer and intra-class reliability was run on the two sets of 15 scores.

#### 2.1.5 Data Analysis

All statistical analyses were conducted using SPSS v 24.0 using a *p* < .05. For the repeated-measures factors, the results were considered significant only if also significant when using the Greenhouse-Geisser correction for any violation of sphericity. For consumption data (sucrose solution intake, chow intake) and body weights, data from Phase 1 and Phase 2 were analyzed separately using mixed ANOVAs. Sucrose solution intakes on common sucrose-access days were analyzed separately for each phase with mixed ANOVAs.

For several analyses of behavioral data, two planned orthogonal contrasts were carried out: (1) between the Chow and the two sucrose groups, and (2) between the Binge and Unrestricted groups.

### 2.2. Results

#### 2.2.1. Consumption data

##### 2.2.1.1. Chow and body weight

As suggested by the mean daily chow intakes and body weights shown in Table 2, no group differences in chow intake were detected either during Phase 1 or Phase 2 (*p*s > .10). There was a linear increase in body weight across the experiment (linear trend *p* < .001), but at similar rates between the groups, with no group differences in body weight found either at the end of Phase 1 or end of Phase 2 (*F*s < 1).

**Table 2.**
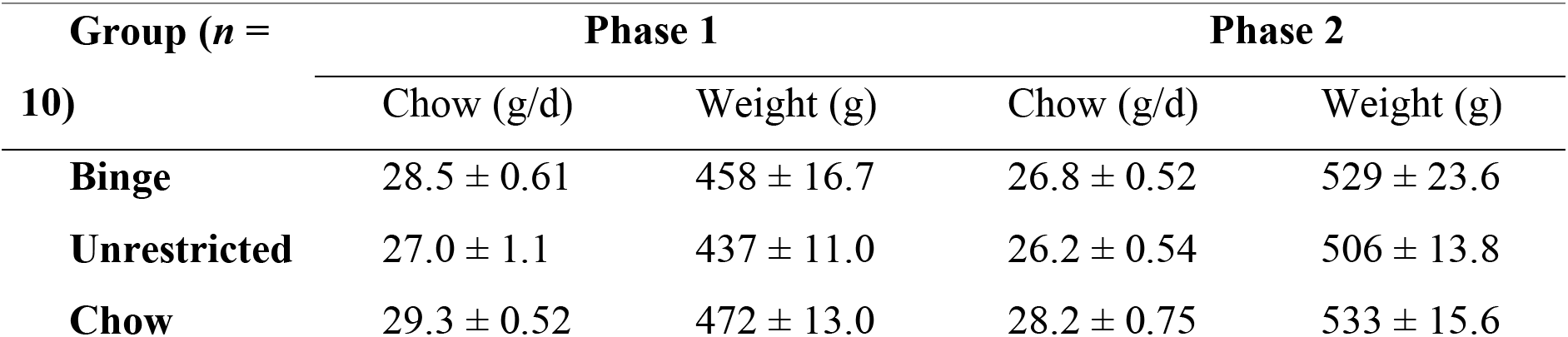
Mean (± SEM) daily chow intake during Phase 1 and 2 and mean (± SEM) body weight at the end of Phase 1 and 2 in Binge, Unrestricted and Chow groups in Experiment 1.

| Group ($n =$<br>10) | Phase 1 | | Phase 2 | |
| --- | --- | --- | --- | --- |
|  | Chow (g/d) | Weight (g) | Chow (g/d) | Weight (g) |
| **Binge** | 28.5 ± 0.61 | 458 ± 16.7 | 26.8 ± 0.52 | 529 ± 23.6 |
| **Unrestricted** | 27.0 ± 1.1 | 437 ± 11.0 | 26.2 ± 0.54 | 506 ± 13.8 |
| **Chow** | 29.3 ± 0.52 | 472 ± 13.0 | 28.2 ± 0.75 | 533 ± 15.6 |

##### 2.2.1.1. Sucrose solution consumption

*Phase 1*. At the beginning of Phase 1, Binge and Unrestricted groups had similar sucrose intakes, *t*(18) = 1.31, *p* > .10 (see Figure 1). Thereafter, intakes remained stable in the Unrestricted group, while by the end of Phase 1the Binge group came to consume more than twice the amount of sucrose in a 23.5-h period than the average for the Unrestricted group. A 3 × (7) Day × Group mixed ANOVA revealed a main effect of Group, *F*(1, 18) = 166.8, *p* < .001, and a linear trend in sucrose intake across days, *F*(1, 18) = 12.36, *p* = .002. This linear trend interacted with Group and Day, *F*(1, 18) = 8.24, *p* = .01. To clarify the nature of the interaction, separate trend analyses were conducted for the Binge and Unrestricted groups. The Binge group showed a linear trend in sucrose intake, *F*(1, 9) = 11.82, *p* = .007, which was not found in the Unrestricted group, *F* < 1, confirming that the Binge group escalated their sucrose intake across Phase 1, whereas the Unrestricted group did not.

**Figure 1.**
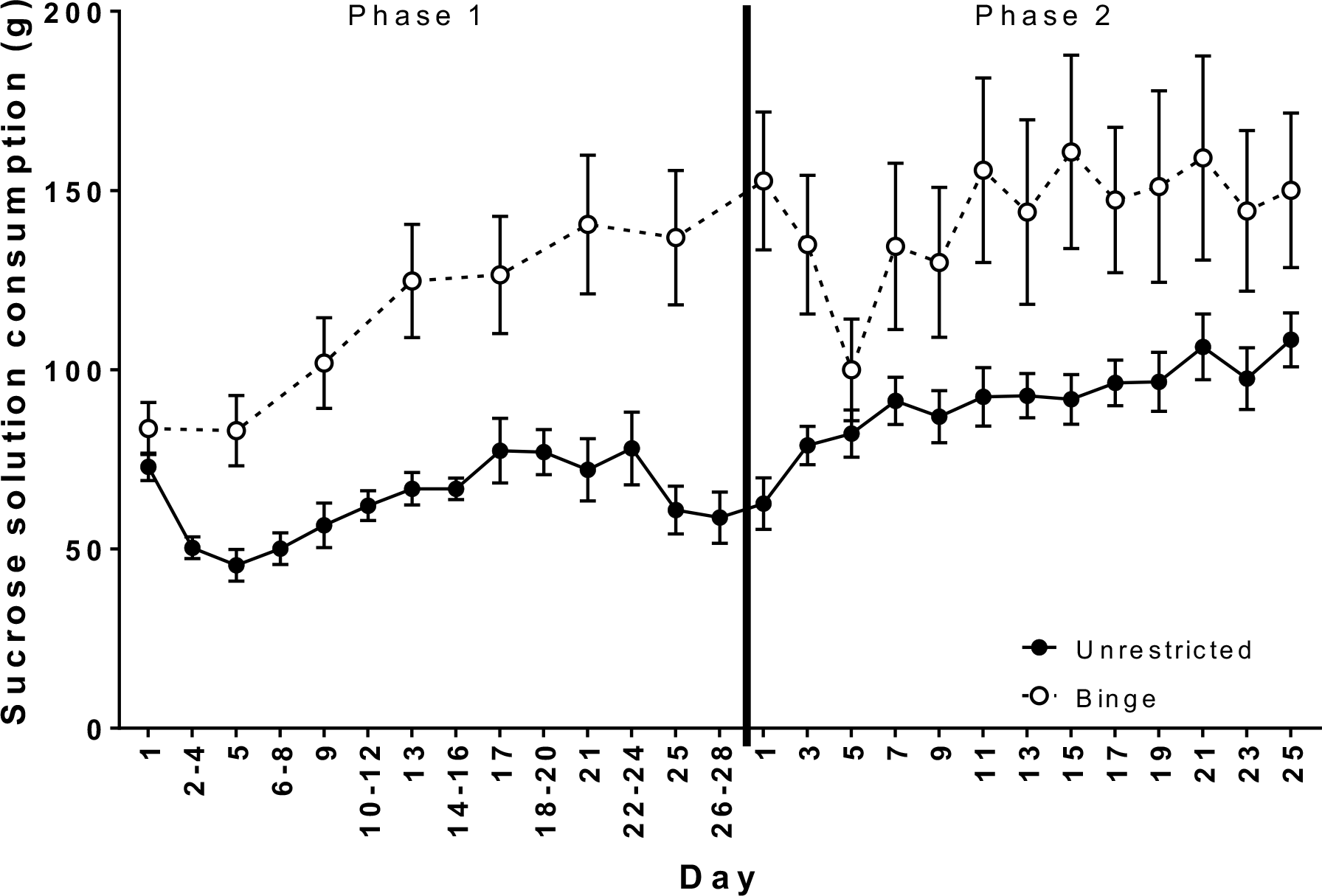
Mean (± SEM) 4% sucrose solution intake in rats given 23.5-h access either every fourth day (Binge) or daily (Unrestricted) in Phase 1. In Phase 2, both groups were given 4% sucrose every second day. Sucrose intakes shown are the amount of sucrose solution consumed in a 23.5-h period. NB: days labelled in this figure correspond to day of the respective phase and not the experimental day.

*Phase 2*. Sucrose intakes in the Binge group decreased over the first three days of Phase 2, before returning to the elevated sucrose intakes found at the end of Phase 1 (see Figure 1). The Unrestricted group increased their sucrose intake across Phase 2 but continued to maintain lower intakes than the Binge group throughout the 28 days of alternate-day access. The 2 × (13) Group × Day mixed ANOVA revealed a significant linear trend in sucrose intake across days, *F*(1, 18) = 21.75, *p* < .001, a Group, *F*(1, 18) = 5.62, *p* = .03 and Group by Day interaction effect, *F*(12, 216) = 2.72, *p* = .03.

As the Binge group displayed a transient decrease in sucrose intakes at the beginning of Phase 2, separate analyses were carried out for the first and last halves of this phase to assess the eventual stability of group intake differences. A 2 × (6) Group × Day mixed ANOVA was conducted for the first six access days and a 2 × (7) Group × Day mixed ANOVA was conducted for the last seven access days of Phase 2. The 2 × (6) mixed ANOVA revealed Group, *F*(1, 18) = 6.76, *p* = .02 and interaction effects, *F*(5, 90) = 6.45, *p* = .004. There were significant linear and quadratic trends in sucrose intake across groups, and an interaction in quadratic trend, *F*(1, 18) = 18.33, *p* < .001. The 2 × (7) mixed ANOVA confirmed that sucrose intake across the last seven days did not significantly increase across days, averaged across groups (*p* > .10) nor was there a Group-by-Day interaction, *F* < 1. Averaged over these last seven days, sucrose intake was significantly higher in the Binge group (*M* = 151.0) than the Unrestricted group (*M* = 98.5), confirming that the Binge group maintained elevated sucrose intakes relative to the Unrestricted group in the second half of Phase 2.

#### 2.2.2. Behavioral data

In summary, behavioral tests of ‘craving’ and ‘withdrawal’ did not yield any differences between Unrestricted and Binge groups, despite the binge-like sucrose consumption exhibited by the latter.

##### 2.2.2.1 Lever-press responding

The mean number of lever presses during the two VR-5 sessions in the Pre-diet Phase was taken as the measure of baseline responding, while response rates at the end of Phases 1 and 2 were based on a single VR-5 session. These data analyzed using a 3 × (3) Group × Test mixed ANOVA which revealed a significant Test effect, *F*(2, 42) = 10.22, *p* < .001, but no effect of Group, *p* > .10, while the Test by Group interaction only approached significance, *F*(4, 42) = 2.13, *p* = .09. Planned contrasts revealed that at the end of Phase 1, lever-press responding was significantly higher in the Chow group (*M* = 144.00) than in the Binge and Unrestricted groups on average, (*M* = 72.69), *F*(1, 21) = 4.97, *p* = .037; however, no difference between Binge and Unrestricted groups was found, *F* < 1. Remaining planned contrasts failed to find differences between sucrose and Chow groups, and between Binge and Unrestricted groups at baseline and the end of Phase 2, all *p*s > .10.

##### 2.2.2.2 Almond preference

Almond preferences are shown in Figure 2A. A 3 × (3) Group × Test mixed ANOVA revealed a significant effect of Group, *F*(2, 27) = 8.10, *p* = .002. No other main effects or interactions were found, *p*s > .10. Planned contrast analyses failed to find any difference in the groups’ almond preferences at baseline, *F* < 1. At the end of Phase 1, almond preferences were significantly higher in the sucrose groups (Binge and Unrestricted on average, *M* = 68.02%) than the Chow group (*M* = 55.41%), *F*(1, 27) = 4.48, *p* =.04; however, no difference between the Binge and Unrestricted groups was detected, *F* < 1. Similarly at the end of Phase 2, almond preferences were higher in the sucrose groups (Binge and Unrestricted on average, *M* = 72.03%) than in the Chow group (*M* = 51.83%), *F*(1, 27) = 10.29, *p* = .003, but again no difference between the Binge and Unrestricted groups was detected, *F* < 1.

**Figure 2.**
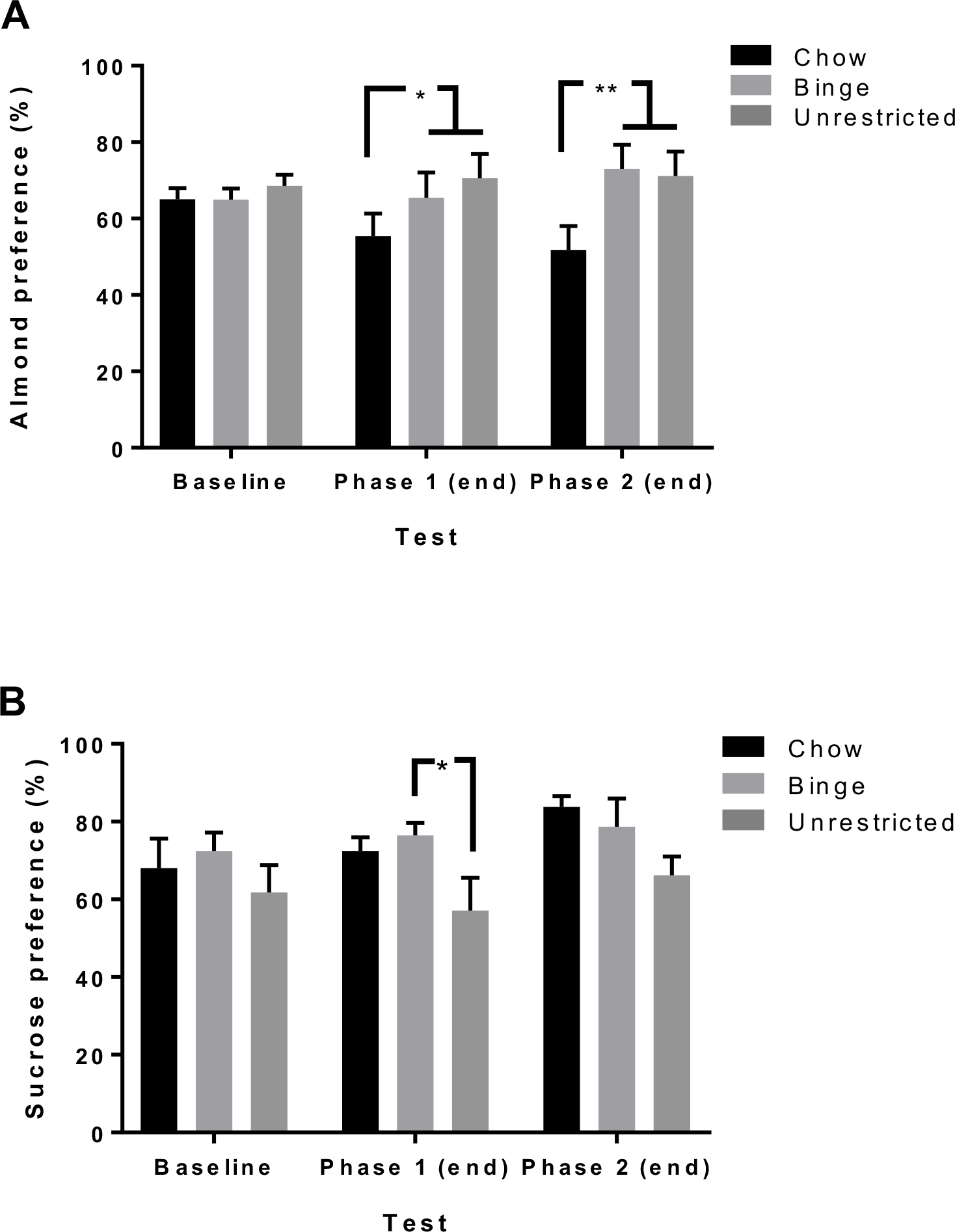
Behavioral data for Experiment 1. A) Mean (± SEM) almond preference in rats at baseline, end of Phase 1 and end of Phase 2. Almond preference was elevated in sucrose groups (Binge and Unrestricted) relative to the Chow group at the end of Phase 1 and 2 (*p*s > .05). B) Mean (± SEM) preference for sucrose over maltodextrin in rats measured at baseline, end of Phase 1 and end of Phase 2. The Binge group displayed higher sucrose preference than the Unrestricted group at the end of Phase 1 (*p* > .05) but this difference disappeared at the end of Phase 2, *p* > .10. * *p* < .05 ** *p* < .01.

##### 2.2.2.3 Sucrose preference

Mean sucrose preference data are shown in Figure 2B. A 3 × (3) Group × Test mixed ANOVA on sucrose preference revealed a significant Group effect, *F*(2, 27) = 3.86, *p* = .03. There was no Test effect or interaction, *Fs* < 1. At baseline, planned contrasts failed to find differences in sucrose preference between sucrose and Chow groups, *F* < 1, nor were there differences between Binge and Unrestricted groups, *p* > .10. At the end of Phase 1, sucrose preference did not significantly differ between sucrose and Chow groups, *F* < 1. However, the Binge group showed significantly higher sucrose preference than the Unrestricted group, *F*(1, 27) = 5.96, *p* = .02. At the end of Phase 2, sucrose preference did not differ between sucrose and Chow groups, nor were there differences between Binge and Unrestricted groups, *p*s > .05.

##### 2.2.2.4 Elevated plus-maze (EPM)

The intra-class correlation coefficient was .99, *p* < .001, indicating high inter-rater reliability. A one-way ANOVA failed to find group differences in the time spent on the open arms of the maze as a percentage of total time spent on the arms, *p* > .10, suggesting that groups demonstrated similar levels of anxiety on the EPM following 48-h sucrose deprivation. The mean percentage of open arm time was 16% in the Chow group, 26% in the Binge group, and 16% in the Unrestricted group.

### 2.3. Discussion

Experiment 1 successfully replicated the persistent elevations in sucrose consumption found in Eikelboom and Hewitt (2016)’s study using a modification of their protocol, whereby the length of Phase 1 was reduced from their 49 days (Eikelboom & Hewitt, 2016; Experiment 1) to the present 28 days. In the Binge group intake of 4% sucrose during Phase 1 increased to almost three times the daily intake of the Unrestricted group, and this difference in intake was still evident after the 28 days of Phase 2. The elevated intakes exhibited by the Binge group satisfied the criteria for binge-like consumption; intakes gradually escalated during Phase 1 and 23.5-h intakes were larger in the Binge group relative to the Unrestricted group under identical access conditions in Phase 2. It may be noted, however, that the absolute amounts of sucrose solution consumed by the Binge group did not reach the level reported by Eikelboom and Hewitt; whereas their Binge group reached a mean of around 300 ml per day after 28 days, the present Binge group reached only 150 ml per day.

A novel feature of this experiment was to add behavioral measures of ‘withdrawal’ and ‘craving’ to the Eikelboom protocol. Although such measures in previous studies have suggested a relationship between sugar bingeing and addiction-like behavior (Avena, Rada, et al., 2008), no such evidence was found in the present experiment. The Binge and Unrestricted groups displayed similar anxiety-like behavior after a 48-h withdrawal period from sucrose, similar motivation to obtain sucrose, and similar preferences for a sucrose-associated flavor. The Binge group only differed from the Unrestricted group in their higher preference for sucrose over maltodextrin at the end of Phase 1 but this difference disappeared by the end of Phase 2. This suggests that intermittent access during Phase 1 may have produced a transient increase in the hedonic value of sucrose. but cannot account for the persistence of binge-like sucrose consumption in the Binge group. It is possible that this experiment failed to find group differences in ‘craving’ because these measures were administered during the diet-intervention instead of sugar withdrawal, as conducted in other studies (Avena et al., 2005). The Chow group may have been more motivated to obtain sucrose than the Binge and Unrestricted groups because they did not receive sucrose in their home cages.

## 3. Experiment 2: Taste or caloric intake?

The main purpose of Experiment 2 was to determine the relative importance of the taste, i.e. sweetness, and of the caloric value of sucrose in producing the persistent bingeing effect demonstrated in Experiment 1. Eikelboom and Hewitt (2016) concluded from their third experiment that intermittent access delays satiety signals. In their experiment, a lick-by-lick analysis of sucrose consumption revealed that the intermittent group had consistently larger sucrose meals compared to the continuous group, but both groups had similar meal initiations. Thus, it appears that the intermittent group engaged in binge-like consumption because they required larger amounts to reach satiety. This suggests then that the caloric value, rather than the taste, of sucrose is a greater driving force behind the persistent bingeing effect.

The basic method used in this second experiment was similar to that used in Experiment 1. The most important changes were to replace 4% sucrose with an isohedonic 0.4% saccharin (non-caloric sweetener) (Young & Trafton, 1964) in two groups (Saccharin Unrestricted; SU, Saccharin Binge; SB), and with an isocaloric 4% maltodextrin (non-sweet, caloric polysaccharide) solution in two further groups (Maltodextrin Unrestricted; MU, Maltodextrin Binge; MB). As detailed below, following a collapse of the bingeing effect in Phase 2, the design was modified to include a third phase (see Table 2).

Experiment 2 used similar behavioral measures of withdrawal and craving to those in Experiment 1: 1) lever pressing on a VR reinforcement schedule; 2) preference for a maltodextrin- or saccharin-paired flavor (i.e. almond) and; 3) preference for maltodextrin or saccharin over an equally attractive sucrose solution. However, in the present experiment post-tests for these measures were conducted after a 7-day withdrawal period following the diet-intervention. As in Experiment 1, possible withdrawal-induced anxiety was measured on the EPM.

It was predicted that during Phase 1, the two Binge groups receiving every-fourth-day access to saccharin (SB) or maltodextrin (MB) would escalate their daily intakes relative to their Unrestricted counterparts (SU and MU groups). Of particular interest was whether elevated consumption in the SB and MB groups would persist during Phase 2, when, as in Experiment 1, both Binge and Unrestricted groups were transferred to the same alternate-day schedule.

### 3.1. Methods

#### 3.1.1. Animals

Forty experimentally-naïve male Sprague-Dawley rats from the same source as arrival and were initially group-housed (*n* = 5/cage). As previously, the temperature- and humidity-controlled colony room was maintained on a reversed 12-h light/dark cycle (lights off at 0900 hrs). After two days of acclimatization and handling, the Pre-diet Phase began with chow and water restrictions identical to those described for Experiment 1. Other details were the same as described for Experiment 1.

Experiment 1 were six weeks old, with an average weight of 308 g (range 285-330 g), on The timeline of Experiment 2 is outlined in Table 3.

**Table 3.**
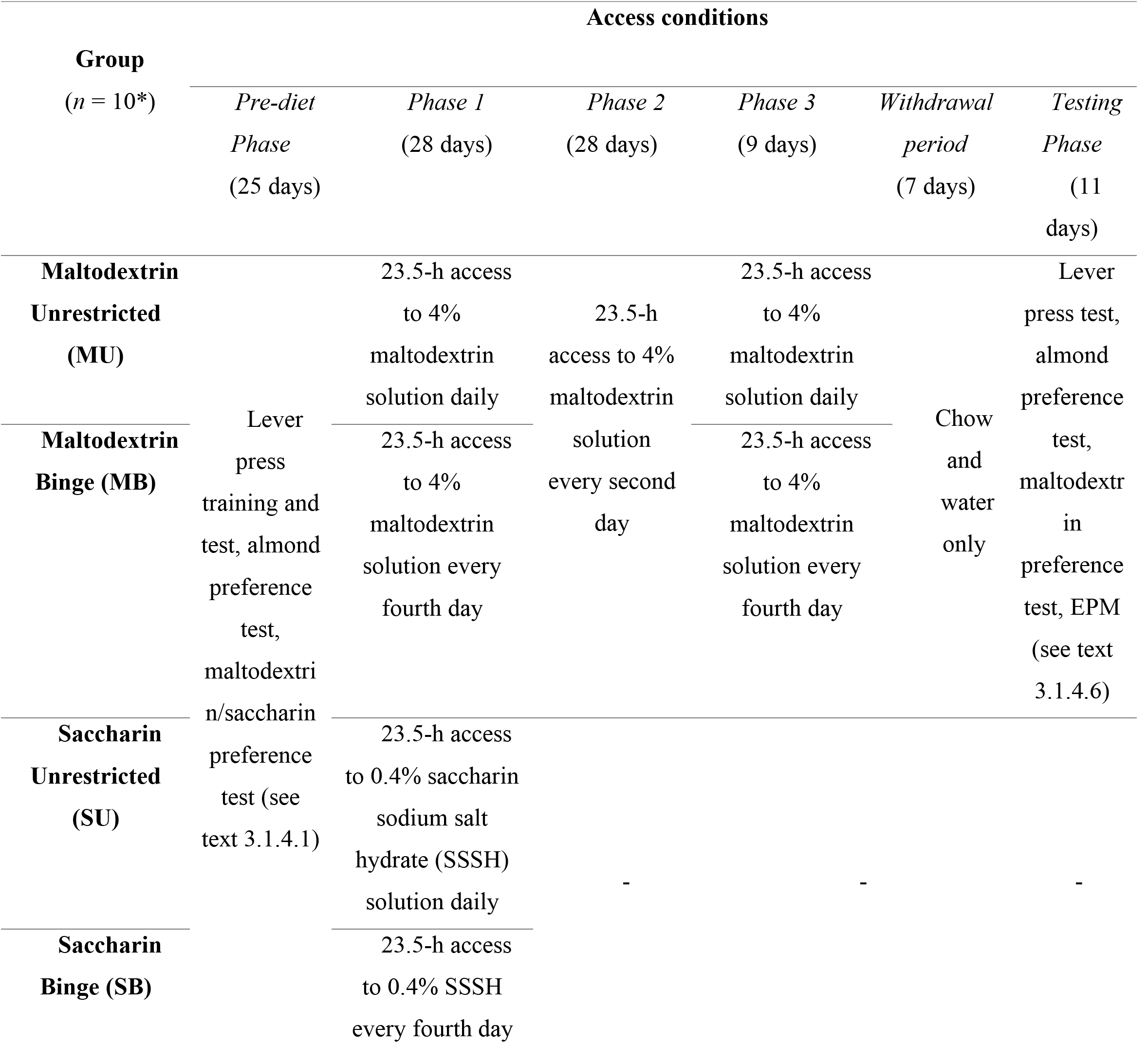
Experimental design of Experiment 2. *Maltodextrin Binge *n = 9* in Phase 2 and 3.

#### 3.1.2. Solutions

Sucrose and maltodextrin solutions were prepared as described for Experiment 1. The 0.4% (w/v) saccharin sodium solution was prepared using saccharin sodium salt hydrate (SSSH; Sigma-Aldrich, S-1002) in the Pre-diet Phase and the majority of Phase 1. Due to a shortage of SSSH in the laboratory at the end of Phase 1, there was an unplanned switch to a ~0.4% (w/v) pure (acid-free) saccharin solution (Sigma-Aldrich, 240931).

#### 3.1.3. Apparatus

The apparatus was identical to that used in Experiment 1.

#### 3.1.4. Procedure

##### 3.1.4.1. Pre-diet phase (Days 1-25)

Rats were first allocated to two weight-matched conditions (*n* = 20/condition) and received either saccharin or maltodextrin solutions throughout the experiment.

*Lever press training (Days 1-24)*. Rats were initially reinforced using a 10% (w/v) sucrose solution. Criteria and procedures for successful magazine training and lever-press training were identical to those in Experiment 1. Up to 18 sessions were given, with data from rats still failing training criterion excluded from consequent analyses (*n* = 5). Rats then received two sessions of VI-10s lever-press training across two days. The first session used a 10% sucrose solution reinforcer. The second session used 4% (w/v) maltodextrin solution as the reinforcer for rats in the Maltodextrin condition, and a 0.4% saccharin solution reinforcer for rats in the Saccharin condition. Rats then received VR-5 lever-press sessions on two successive days.

*Almond preference training and test (Days 3-6)*. The procedure was identical to that of Experiment 1, except that the Maltodextrin rats were trained using a 4% (w/v) maltodextrin + 1% (v/v) almond solution and the Saccharin rats were trained using a 0.4% SSSH (w/v) + 1% (v/v) almond solution. During the two-bottle choice tests, the base solution in the Maltodextrin condition was 1% (w/v) maltodextrin, and in the Saccharin condition was 0.1% (w/v) SSSH solution. Each rat was given a two-bottle choice test between almond + base and base only. Other details were the same as for Experiment 1.

*Target solution preference test (Day 8)*. Each rat received a preference test using the procedure described in Experiment 1. The Maltodextrin rats were given a two-bottle choice test between 4% maltodextrin and 4% sucrose, while those in the Saccharin condition were given a two-bottle choice test between 0.4% SSSH and 2% sucrose. The choice of a 2% sucrose solution in the latter test was to avoid a possible floor effect, since pilot tests had indicated that comparison with a 4% sucrose solution produced a low saccharin preference (~23%). In the first half of each test session, maltodextrin or saccharin was placed on the left and sucrose was placed on the right. Maltodextrin preference was calculated as the consumption of 4% maltodextrin solution as a percentage of total fluid consumption (4% maltodextrin and 4% sucrose) in the two-bottle choice tests. Saccharin preference was calculated as the consumption of 0.4% saccharin solution as a percentage of total fluid consumption (0.4% saccharin and 2% sucrose) in the two-bottle choice tests.

##### 3.1.4.2. Phase 1 (Days 26-53)

At the start of this phase rats in the Maltodextrin condition were allocated to two groups (*n* = 10/group) matched primarily for body weight and almond preference but also to a lesser degree for sucrose preference and lever-press responding. One group was labeled the Maltodextrin Binge (MB) group; this was given 23.5-h access to 4% maltodextrin solution every fourth day, starting at 1000 hrs and ending at 0930 hrs the next day. The other was labeled the Maltodextrin Unrestricted (MU) group; this received 23.5-h access to 4% maltodextrin solution daily.

Rats in the Saccharin condition were similarly allocated to two matched groups (*n* = 10/group). The Saccharin Binge (SB) group received 23.5-h access to 0.4% SSSH solution every fourth day, starting at 1000 hrs and ending at 0930 hrs the next day, while the Saccharin Unrestricted (SU) group received 23.5-h access to 0.4% SSSH solution daily. Due to an unavailability of SSSH at the end of Phase 1, there was an unplanned switch to pure saccharin solutions. This caused an abrupt decrease in saccharin intakes in the SU group on the last 3 days of Phase 1 (see Figure 3). Consequently, only Phase 1 data will be reported here for the SB and SU groups.

**Figure 3.**
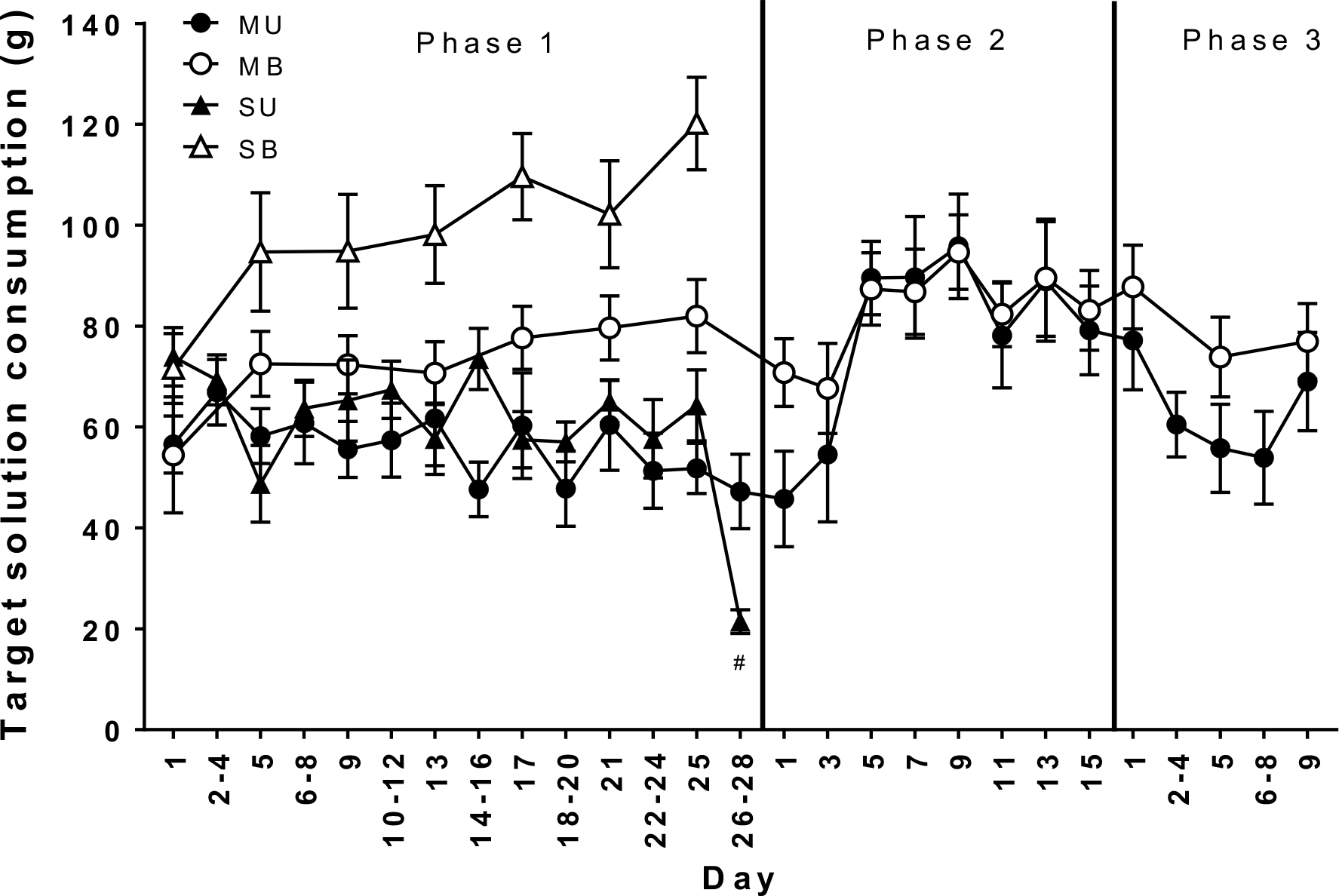
Mean (± SEM) 4% maltodextrin (MU, MB) or 0.4% saccharin (SU, SB) solution intakes in a 23.5-h period in rats given 23.5-h access every fourth day (MB, SB) or daily (MU, SU) in Phase 1. In Phase 2, the MB and MU groups were given 23.5-h access to 4% maltodextrin solution every second day. In Phase 3, MB and MU groups were returned to their respective Phase 1 access schedules. Both Binge groups (MB, SB) drank significantly greater amounts by the end of Phase 1 relative to their Unrestricted counterparts (MU, MB), *p* < .001. The intake difference between MU and MB groups disappeared in Phase 2 (*F* < 1), and was not reinstated in Phase 3 (*p* > .10). # indicates that this drop in intakes followed an unplanned switch from SSSH solution to pure saccharin solution in SU group; the SU and SB groups were dropped from Experiment 2 because of SSSH unavailability and subsequent data have not been reported. NB: Days in this figure indicate the day of each respective phase and do not correspond to the experimental day.

##### 3.1.4.3. Phase 2 (Days 54-69): MB and MU groups only

The MB and MU groups were switched to an alternate-day access schedule such that they received 23.5-h access to 4% maltodextrin every second day from 1000 hrs to 0930 hrs the next day.

##### 3.1.4.4. Phase 3 (Days 70-78): MB and MU groups only

As detailed below (see Section 3.2.1.1), the intake difference between the MB and MU groups seen in Phase 1 collapsed during Phase 2. Consequently, a third phase was added in which the groups were switched back to their Phase 1 conditions for nine days in an attempt to reinstate binge-like consumption. Thus, the MB group was again given 23.5-h access to 4% maltodextrin every fourth day, whereas the MU group was again given 23.5-h access to 4% maltodextrin daily.

##### 3.1.4.5. Withdrawal period (Days 79-85): MB and MU groups only

During this one-week period no further access to maltodextrin solution was given and rats were maintained on *ad libitum* chow and water only.

##### 3.1.4.6. Testing phase (Day 86-96): MB and MU groups only

On Day 85, chow was removed at 1700 hrs to mimic the mild food deprivation induced during baseline tests in the Pre-diet Phase. From Day 86-88 each rat was given three lever-press sessions over three successive days (one session/day), using the following reinforcement schedules: VI-10s, VR-5, VR-5. The average number of lever presses over the two VR-5 sessions was taken as the after-withdrawal measure of lever-press responding. On Day 91 rats were tested for almond preference using the two-bottle choice test procedure. The next day rats were also tested for preference for maltodextrin over 4% sucrose solution as previously described. During these test days, rats were given 2-h daily chow access at 1400 hrs (after their daily lever-press session or two-bottle choice test). Rats were returned to *ad libitum* chow on Day 93 before EPM testing. On Day 94 each rat was tested on the EPM following the procedure described for Experiment 1. Behavior on the EPM was scored and calculated as described for Experiment 1. To establish inter-rater reliability, ten EPM recordings were also rated by a blind scorer and intra-class reliability between the two sets of ten scores was measured.

#### 3.1.5. Data analysis

One rat from the MB group died during Phase 2 and its data were excluded from all analyses subsequent to Phase 1. As detailed earlier, only target solution intake data from Phase 1 was analyzed for the SB and SU groups and no other intake or behavioral data are reported for these groups. Due to the change in access conditions between phases, target solution intakes on common access days were analyzed separately for each phase with mixed ANOVAs. Since motivational states may have differed across tests of lever-pressing and flavor preferences due to differences in the degree of food deprivation, overall changes across tests were not considered meaningful and consequently only data from after-withdrawal tests were analyzed using one-way ANOVAs.

### 3.2. Results

#### 3.2.1. Pre-diet measures

Although the concentrations of saccharin and maltodextrin in the present experiment were chosen to match the hedonic and caloric value of 4% sucrose, data from the pre-diet measures suggested that 0.4% saccharin was a more effective reinforcer than 4% maltodextrin. During lever-press training and baseline test where rats were reinforced with their target solution i.e. 0.4% saccharin or 4% maltodextrin, rats in the Saccharin condition had higher average lever-press responding on a VR-5 reinforcement schedule than rats in the Maltodextrin condition (148 vs. 87), however this only approached significance (*p* = .05).

#### 3.2.2. Consumption data

##### 3.2.2.1. Chow and body weight

As suggested by Table 4, there were no main effects on chow intake but there was a significant Solution × Access interaction (*F*(1, 36) = 87.0, *p* < .001) such that the MU group had lower chow intakes than the MB group, whereas no difference in chow intakes was detected between the SU and SB groups. This suggests that the MU group reduced chow intake to compensate for the additional caloric intake from unrestricted access to a caloric maltodextrin solution, whereas this was unnecessary for the SU group because the saccharin solution contained no calories. In Phase 2 and Phase 3 there were no differences in daily chow intake between MU and MB groups (*p* > .05). Body weights gradually increased across the experiment, but no group differences were found (*p*s > .10).

**Table 4.** Mean (± SEM) daily chow intake during Phases 1-3, and mean (± SEM) body weight at the end of Phase 1-3 in MU and MB groups in Experiment 2. Data for SU and SB groups are shown only for Phase 1 (see section 3.1.4.2 for details). *Maltodextrin Binge *n = 9* in Phase 2 and 3.

| Group ($n = 10^*$) | Phase 1 | | Phase 2 | | Phase 3 | |
| --- | --- | --- | --- | --- | --- | --- |
|  | Chow<br>(g/d) | Weight<br>(g) | Chow<br>(g/d) | Weight<br>(g) | Chow<br>(g/d) | Weight<br>(g) |
| **Maltodextrin Unrestricted (MU)** | 25.6 ±<br>0.68 | 472 ±<br>16.1 | 24.1 ±<br>0.57 | 507 ±<br>21.2 | 24.5 ±<br>0.49 | 528 ±<br>22.8 |
| **Maltodextrin Binge (MB)** | 28.9 ±<br>0.36 | 490 ±<br>14.8 | 26.8 ±<br>0.59 | 522 ±<br>20.4 | 27.7 ±<br>0.28 | 542 ±<br>21.3 |
| **Saccharin Unrestricted (SU)** | 29.3 ±<br>0.68 | 483 ±<br>15.3 | - | - | - | - |
| **Saccharin Binge (SB)** | 28.1 ±<br>0.61 | 467 ±<br>15.0 | - | - | - | - |

##### 3.2.2.2. Target solution consumption

*Phase 1*. Mean daily intakes of target solutions on access days are shown in Figure 3. No group differences were found in solution intakes on Day 1 of Phase 1, *p* > .10. Subsequently, MB and SB groups came to consume increasing amounts of their target solution, whereas no increases in the daily intakes by the MU and SU groups were found. Intakes of saccharin were generally greater than intakes of maltodextrin. However, saccharin intakes dropped due to a switch in saccharin solutions on Day 26 of Phase 1. This description of the results was confirmed by a 2 × 2 × (7) Solution × Access × Day mixed ANOVA that revealed a main effect of Day, *F*(6, 216) = 4.70, *p* < .001, and a Day by Access interaction effect *F*(6, 216) = 8.35, *p* < .001. There were main effects of Solution (maltodextrin vs. saccharin), *F*(1,36) = 5.33, *p* = .027 and of Access (unrestricted vs. binge), *F*(1,36) = 16.01, *p* < .001, revealing that SU and SB groups had higher intakes than the MU and MB groups (80.25g vs. 65.27g, on average), and that MB and SB groups drank considerably more than MU and SU groups (85.75g vs. 59.78g, on average), averaged over solution.

There was a significant linear trend of intake across days, *F*(1,36) = 16.90, *p* < .001. This trend interacted with Access (*F*(1, 336) = 19.30, *p* < .001), suggesting that the pattern of intake across days differed between groups receiving unrestricted access (SU, MU) and those receiving binge access (SB, MB). To clarify the nature of the interaction, separate trend analyses were conducted for access conditions. For unrestricted-access groups (MU and SU), there was no significant trend in intakes across days, *p*s > .10. For binge-access groups (MB and SB), there was a significant linear trend in intake, *F*(1, 18) = 28.44, *p* < .001.

The Solution by Access interaction was not significant, *p* > .10, indicating that the degree to which the binge condition elevated intakes above those in the unrestricted condition was not detectably different between maltodextrin and saccharin. On average MB rats came to consume 1.5 times the amount of maltodextrin relative to the MU rats, whereas SB rats came to consume almost twice the amount of saccharin relative to SU rats.

*Phase 2*. As seen in Figure 3, within three alternative-day exposures in Phase 2 there were no longer any differences in intakes between the MB and MU groups and they remained similar until the end of the phase. A 2 × (8) Access × Day mixed ANOVA revealed significant linear, *F*(1, 17) = 32.23, *p* < .001 and quadratic, *F*(1, 17) = 39.83, *p* < .001 trends in maltodextrin intake across days. There were also significant interactions in linear, *F*(1, 17) = 6.29, *p* = .023, and quadratic, *F*(1, 17) = 7.93, *p* = .012, trends between Day and Access. There was no Access effect (*F* < 1), which confirms that the MB and MU groups did not differ in terms of maltodextrin intakes in Phase 2.

*Phase 3*. The results shown for Phase 3 of Figure 3 suggest that MU and SU rats decreased their intakes upon reinstatement of unrestricted access, while MB and SB rats maintained higher intakes when given every-fourth-day access. However, a 2 × (3) Group × Day mixed ANOVA revealed only linear, *F*(1, 17) = 5.52, *p* = .031, and quadratic trends, *F*(1,17) = 20.96, *p* < .001, but no other main effects or interactions, *p*s > .10. Thus, binge-like consumption in the MB group relative to the MU group was not reinstated in Phase 3.

#### 3.2.3. Behavioral data

In summary, no differences were found between the MB and MU group on any of the behavioral measures. A one-way ANOVA applied to lever-press rates did not find any Group effect (*F* <1), indicating that both MU and MB groups exhibited similar lever-press responding after withdrawal. Mean almond preference data are shown in Figure 4A. A one-way ANOVA on almond preference after withdrawal confirmed that there was no difference between the MU and MB groups (*F* < 1). Mean target solution preference data are shown in Figure 4B. As seen in this figure, the preference for maltodextrin over sucrose was similar after withdrawal in the MU and MB, *p* > .10. When videos of performance on the elevated plus maze were scored, the intra-class correlation coefficient was .99, *p* < .001, indicating very high inter-rater reliability. Average open-arm time was 10.3% in the MU group and 14.6% in the MB group. A one-way ANOVA failed to find differences between the MU and MB groups in the percentage of time spent on the open arms of the maze (*F* < 1), suggesting that MB and MU rats exhibited similar levels of anxiety-like behavior. It should be noted that these low open-arm times suggest that both groups were relatively anxious.

**Figure 4.**
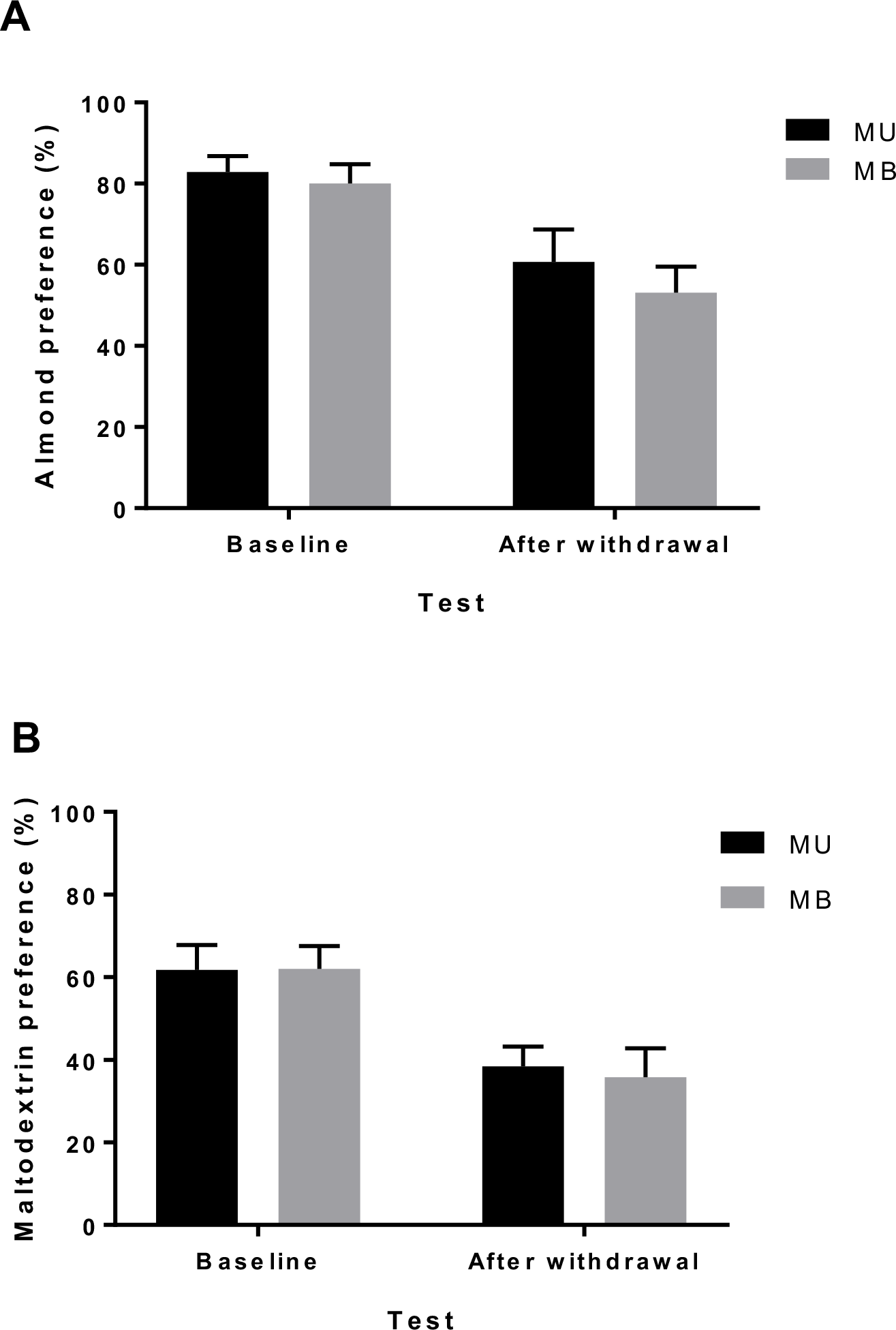
Behavioral data for Experiment 2. A) Mean (± SEM) almond preference in at baseline and after withdrawal. No difference in almond preference was found between MB and MU groups (*F* < 1). B) Mean (± SEM) preference for 4% maltodextrin solution over 4% sucrose solution at baseline and after withdrawal in MU and MB rats. Maltodextrin preference was similar in MB and MU groups (*p* > .10).

### 3.3. Discussion

As with sucrose in Experiment 1, in Phase 1 of the present experiment providing access only every fourth day produced increasing intakes of both the maltodextrin and saccharin solutions, while in the Unrestricted condition intakes of these solutions showed little change. In contrast to the results for sucrose found in Phase 2 of Experiment 1, in the present experiment differences in intake between the Binge group given maltodextrin (MB) and the group given unrestricted access to maltodextrin (MU) were not maintained in Phase 2. Furthermore, in Phase 3 returning the two groups to their conditions in Phase 1 failed to re-instate the previous differences.

The concentration of the maltodextrin solution in the present experiment was chosen to match the energy content of the 4% sucrose solution used in Experiment 1. Consequently, the failure to find persistent binge-like consumption of maltodextrin suggests that energy content is not an important contributor to this effect but rather that the sweet taste of sucrose plays an important role. Due to the unplanned switch in saccharin solutions, however, we were unable to assess whether the binge effect with a non-caloric sweet solution would persist in the present experiment. Following the present experiment we carried out a systematic comparison between sodium saccharin salt hydrate (SSSH) and pure (acid-free) saccharin (S). This confirmed that 0.4% SSSH is much more acceptable to rats than 0.4% S, whereas this difference is less apparent at concentrations of 0.1% (Rehn, Onuma, Rooney, & Boakes, 2018)

Regarding the failure in this experiment to find any group differences in the craving and withdrawal measures, this is discussed in the General Discussion.

## 4. Experiment 3: Bingeing on highly hedonic solutions

As for Experiment 3, the main aim of this experiment was to test whether the persistence effect that can be obtained with 4% sucrose can also be obtained using other solutions. While Experiment 2 failed to obtain the effect with a 4% maltodextrin solution, it left open the possibility that the bingeing on 0.4% saccharin sodium salt hydrate (SSSH) solution found in Phase 1 would persist into Phase 2. Therefore, the present experiment included 0.4% SSSH as one of the target solutions. The other target solution was a mixture of 4% glucose and 0.4% SSSH. This was selected because such mixtures are known to be exceptionally palatable to rats (Valenstein, Cox, & Kakolewski, 1967). Although containing no more energy than 4% sucrose, it has a higher hedonic value, as confirmed in the present experiment.

Experiment 3 employed a 2 × 2 factorial design (see Table 5), in which one factor, Solution, was whether rats were given saccharin or the glucose-saccharin mixture, and the other factor was whether rats had access to their solutions on every fourth day (Binge condition) or Unrestricted access during Phase 1. This design generated four groups: Saccharin Unrestricted (SU), Saccharin Binge (SB), Glucose-Saccharin Unrestricted (GSU), Glucose-Saccharin Binge (GSB). It may be noted that, since the glucose + saccharin solution differed from saccharin alone in both being more palatable and containing more energy, we did not plan to draw any general conclusions regarding the basis of such bingeing from potential differences in the size and persistence of a binge effect produced by the two solutions.

**Table 5.**
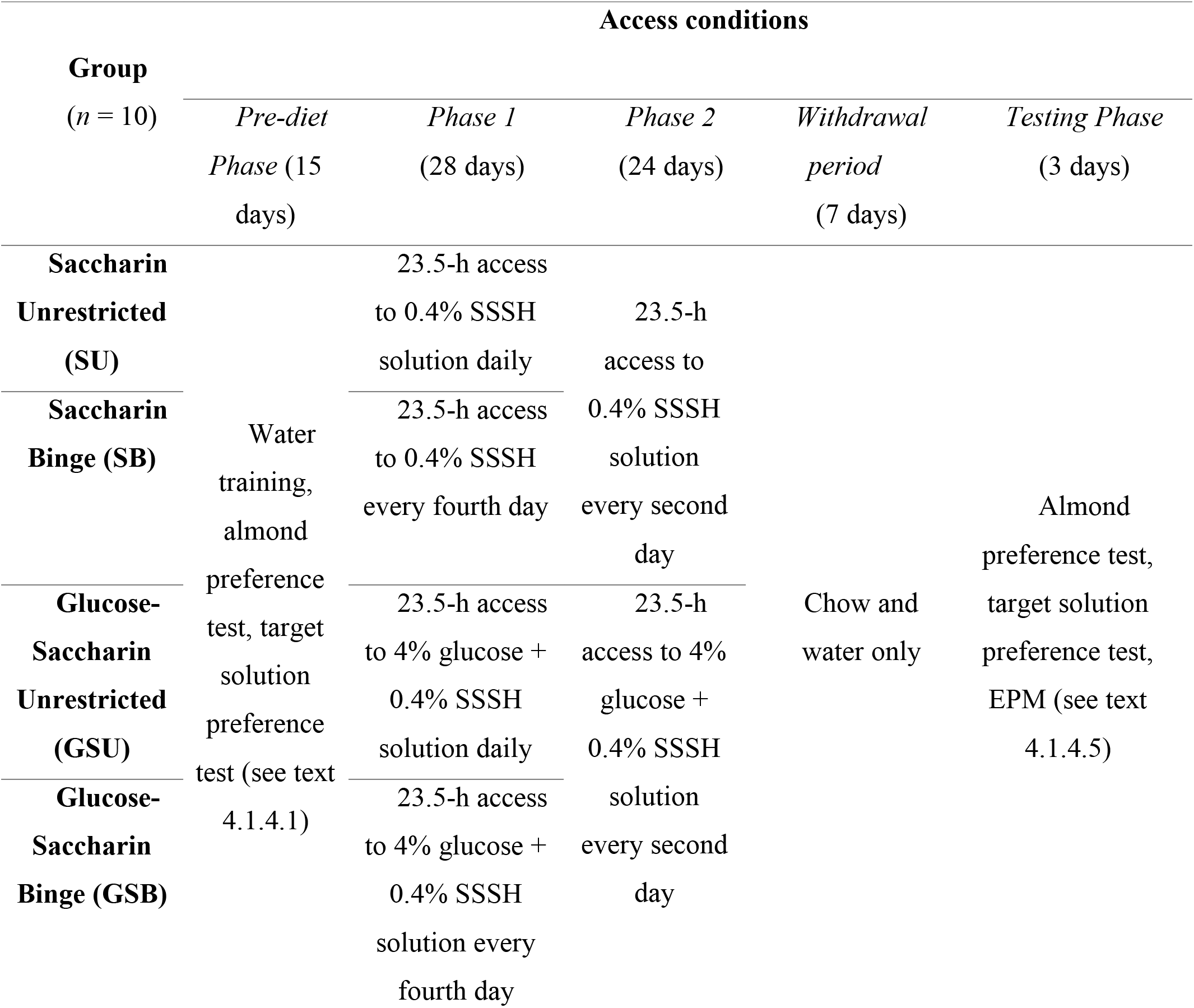
Experimental design of Experiment 3.

The same behavioral measures of ‘withdrawal’ and ‘craving’ used in the previous experiments were employed in Experiment 3, except that lever-press tests were omitted because of the large individual variability in response rates found in Experiments 1 and 2.

### 4.1. Methods

#### 4.1.1. Subjects

Forty experimentally-naïve male Sprague-Dawley rats from the same source as in the previous experiments were eight weeks old, with an average weight of 234 g (range 212 – 281 g), at the start of the experiment. Upon arrival, rats were group-housed (*n* = 4/cage) in a temperature- and humidity-controlled room on a reverse 12:12 h light cycle (lights off at 1000 h). Rats were individually housed in ventilated cages (Techniplast, Australia) divided into two compartments so that an animal had visual, auditory and olfactory contact with its neighbor but no physical contact. Other details are the same as detailed for Experiment 1.

#### 4.1.2. Solutions

As previously, these were prepared in tap water. Saccharin sodium salt hydrate (SSSH; Sigma S-1002) was exclusively used to prepare both the 0.4% saccharin solution and the mixture of 0.4% saccharin and 4% glucose (15.4 kJ/g, Myopure Dextrose Monohydrate (D-glucose) www.myopure.com.au). Sucrose solutions were prepared as described for Experiment 1.

#### 4.1.3. Apparatus

The EPM was at a height of 50 cm above the floor and the video camera was mounted at a height of 92 cm above the center of the EPM. Other details and apparatus are identical to those described for Experiment 1.

#### 4.1.4. Procedure

The timeline of Experiment 3 is outlined in Table 5.

##### 4.1.4.1. Pre-diet Phase (Day 1-15)

After acclimatization and handling, daily water access was gradually reduced across four consecutive days from 4 h, to 2 h, 1 h, and 30 min, in preparation for water training in the drinking chambers. Rats were given 30-min access to water after each water training session.

*Water training (Days 1 – 3)*. Rats were transferred from their home cages to the individual drinking chambers where they were given 30-min access to water in each daily training session. On Day 1 rats were given a single bottle, whereas on Days 2 and 3 rats were given two bottles (both containing water) and the bottle positions were swapped after 15 min. This was done to acclimate rats to the choice test procedure. After the Day 1 session rats were returned to their home cages where half of the rats (*n* = 20) received 4-h access to the 0.4% saccharin solution, while the other half received 4-h access to the 4% glucose + 0.4% saccharin solution. After the Day 2 session rats received 4-h access to the other solution. Rats were subsequently returned to *ad lib* water access.

*Acceptance tests* (Days 4-5). Each rat was given two acceptance tests, one for the saccharin solution and the other for the glucose and saccharin mixture. During these tests a rat received a single bottle of either solution for 30 min in the individual drinking chambers. Rats were then allocated to two conditions (saccharin only vs. saccharin + glucose, *n* = 20/condition) matched for body weight and saccharin acceptance, calculated as 30-min intake. From this point onwards, rats were trained and tested using only their target solutions.

*Almond preference training and test* (Days 6-9). The procedure was essentially the same as that described for the previous experiments. Water bottles were removed from home cages 2 h before testing. Across three consecutive days, each rat underwent one daily 10-min almond preference session where they received either 1% almond + 4% glucose + 0.4% saccharin (saccharin + glucose condition) or 1% almond + 0.4% saccharin (saccharin only condition). These solutions were presented in a single bottle on the first day and presented in two bottles on the second and third days to acclimate rats to the two-bottle choice procedure. After the training, each rat received a 10-min almond preference test where they were presented with 1% almond + base solution in one bottle, and base solution alone in another. The bottle positions were swapped halfway through the test. For the choice tests the base solution was 0.1% saccharin for rats in the saccharin only condition and a mixture of 1% glucose and 0.1% saccharin for rats in the mixed condition.

*Target solution preference tests* (Day 11-15). Water bottles were removed from home cages 2 h before each test. On Day 11 each rat was given a two-bottle choice test between 2% sucrose and their target solution. As all rats showed very high preferences (averaging 83% for saccharin and 93% for the mixture) for their target solution over 2% sucrose, several two-bottle choice tests were conducted using increasing sucrose concentrations (4%, 6%, 8%, 12%) as the comparison solution until preference for the target solution was between 50-70%. The final comparison solution for the saccharin rats was 6% sucrose and the comparison solution for the mixture rats was 12% sucrose. The initial position of the sucrose bottle was counterbalanced within each group.

##### 4.1.4.2. Phase 1 (Day 16-43)

Rats in the saccharin condition were allocated to two groups (*n* = 10/group), Saccharin Unrestricted (SU) and Saccharin Binge (SB), matched for body weight, almond preference and sucrose preference. Rats in the mixture condition were similarly allocated to two matched groups (Glucose + Saccharin Unrestricted [GSU] and Glucose + Saccharin Binge [GSB]). The two Binge groups (SB, GSB) received 23.5-h access to their target solution every fourth day, starting at 1000 hrs and taken off at 0930 hrs the next day, while the two Unrestricted groups (SU, GSU) received 23.5-h access to their target solution daily.

##### 4.1.4.3. Phase 2 (Day 44-67)

All groups were switched to an alternate-day access schedule; rats received access to their target solutions every second day, starting at 1000 hrs and taken off at 0930 hrs the next day. On Day 45 (non-target-solution day) all groups were given an almond preference test using the two-bottle choice test procedure previously described. On Day 47 (non-target-solution day) all groups were given a target solution preference test relative to 6% sucrose for the saccharin groups and relative to 12% sucrose for the mixture groups.

##### 4.1.4.4. Withdrawal period (Day 68-74)

No further access to the target solutions was given for the remainder of the experiment. All remained with unrestricted access to chow and water. For three consecutive days during this period rats were transported in individual transport cages in squads of ten to the EPM testing room for 30 min per day to habituate them to the EPM test procedure.

##### 4.1.4.5. Testing Phase (Day 75-78)

Preference tests were conducted using an identical two-bottle choice procedure to that described in the Pre-diet Phase of this experiment. On Day 75 rats were tested for almond preference. The next day rats received a sucrose preference test. On Days 77 and 78 each rat was tested on the elevated plus-maze (EPM) for 5 min following the procedure described in previous experiments. In this experiment, however, each rat was transferred into individual transport cages and allowed to habituate to the conditions of the EPM testing room for 10 min before being placed on the EPM. To avoid potential testing day effects or time of day effects, the order in which the groups and rats were tested was completely counterbalanced across both days. The videos were scored by a non-blinded experimenter as described for Experiment 1.

#### 4.1.5. Data analysis

As in the previous experiments, consumption data were analyzed separately for Phase 1 and Phase 2 using mixed ANOVAs with Solution and Access as between-subject factors, and Day as the within-subject factor. However, unlike in the previous experiments, Test was included as a factor for the behavioral data in Experiment 3 because rats were sated during both pre- and post-diet tests.

### 4.2. Results

#### 4.2.1. Consumption data

##### 4.2.1.1. Chow and bodyweight

Chow intakes and bodyweights are shown in Table 6. Chow intake in Phase 1 followed a linear trend (*F*(1, 36) = 10.09, *p* = .003) but this did not differ between groups as main effects of Solution or Access or interactions failed to reach significance (all *p*s > .10). Similarly in Phase 2, there was a linear trend in chow intake (*F*(1, 36) = 43.49, *p* < .001) which did not differ between Solution or Access conditions (all *ps* > .10). There was a linear trend in body weight across the experiment (*p* < .001), indicating that all rats gained weight throughout the experiment but this did not differ between groups as no other main effects or interactions were found (all *p*s > .10).

**Table 6.** Mean (± SEM) daily chow intake during Phase 1 and 2 and mean (± SEM) body weight at the end of Phase 1 and 2 in SU, SB, GSU, GSB groups in Experiment 3.

| Group ($n =$<br>10) | Phase 1 | | Phase 2 | |
| --- | --- | --- | --- | --- |
|  | Chow (g/d) | Weight (g) | Chow (g/d) | Weight (g) |
| **Saccharin**<br>**Unrestricted**<br>**(SU)** | 26.8 ± 0.32 | 521 ± 14.4 | 26.6 ± 0.44 | 575 ± 17.1 |
| **Saccharin**<br>**Binge (SB)** | 27.4 ± 0.32 | 534 ± 13.5 | 27.8 ± 0.36 | 600 ± 17.4 |
| **Glucose +**<br>**Saccharin**<br>**Unrestricted**<br>**(GSU)** | 25.7 ± 0.32 | 548 ± 7.8 | 26.2 ± 0.18 | 609 ± 13.2 |
| **Glucose +**<br>**Saccharin**<br>**Binge (GSB)** | 26.2 ± 0.38 | 523 ± 15.1 | 25.5 ± 0.27 | 589 ± 18.3 |

##### 4.2.1.2. Target solution intakes

*Phase 1*. Mean intakes of the target solutions are shown in Figure 5. Initial intakes were high on the first day of Phase 1, but dropped and stabilized around Day 9. A 2 × 2 × (7) Solution × Access × Day mixed ANOVA was conducted on intakes during the days when all rats had access to their target solution. This revealed main effects of Day, *F*(6, 216) = 44.78, *p* < .001, Solution, *F*(1, 36) = 28.80, *p* < .001, Access, *F*(1, 36) = 20.34, *p* < .001, and a Day by Access interaction effect, *F*(6, 216) = 6.759, *p* < .001. As Figure 5 suggests, these main effects reflect: (1) greater intakes in the GSU and GSB groups on average, than the SU and SB groups; (2) greater intakes in the rats that received Binge access (GSB and SB groups) than Unrestricted access (GSU and GSB groups); and (3) a linear, *F*(1, 36) = 55.04, *p* < .001, and quadratic trend, *F*(1, 36) = 82.89, *p* < .001 in target solution intake across days. This analysis also revealed an interaction between Access and linear trend, *F*(1, 36) = 4.32, *p* = .04, indicating that apart from the initial drop in intakes across groups, the elevation in intakes in the GSB and SB groups on average was greater than that seen in the GSU and SU groups. On the other hand, no interaction between Solution and Access (*p* > .10) was detected, indicating that no difference was detected between the saccharin and mixed solutions in the extent to which the one-in-four-days schedule increased intakes above the daily mean intakes by the unrestricted groups.

**Figure 5.**
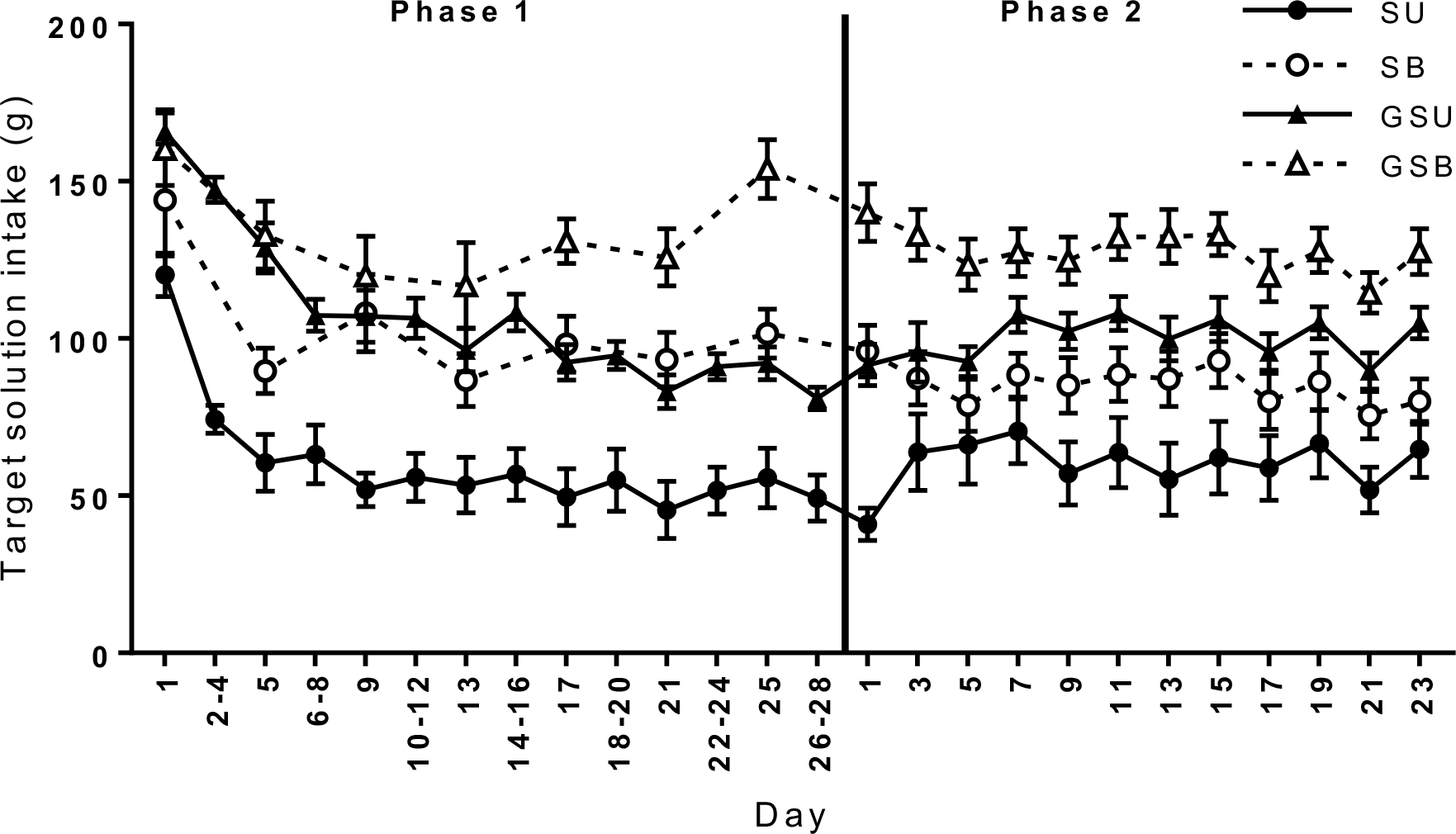
Mean (±SEM) 23.5-h intake of target solution in rats given unrestricted (GSU) or binge access (GSB) to a 4% glucose + 0.4% saccharin solution or unrestricted (SU) or binge access (SB) to a 0.4% saccharin (SSSH) solution. In Phase 1 GSU and SU rats were given daily access to their target solutions, whereas GSB and SB rats were given access every fourth day. In Phase 2 all rats were switched to alternate-day access. SB and GSB groups drank significantly greater amounts of their target solution relative to their unrestricted counterparts by the end of Phase 1 (*p* < .001). This intake difference (i.e. binge effect) was maintained throughout Phase 2 (*p* < .001). NB: Intake data for the GSU group on Day 2-4 is an average of the Day 1 and Day 5 intakes because their bottles were empty upon measurement on Day 4 and intakes would have been higher if they were not limited to the remaining amount of solution in the bottle. Days in this figure indicate the day of each respective phase and do not correspond to the experimental day.

*Phase 2*. As shown in Figure 5, the GSU and SU groups gradually increased their intakes across Phase 2 in the GSU and SU groups and yet the GSB and SB groups maintained consistently elevated intakes. This description was confirmed by a 2 × 2 × (12) mixed ANOVA applied to common target solution-access days in Phase 2. This revealed a main effect of Day, *F*(11, 396) = 5.13, *p* < .001, Solution, *F*(1, 36) = 31.72, *p* < .001, Access, *F*(1, 36) = 13.40, *p* = .001, and a Day by Access interaction, *F*(11, 396) = 4.88, *p* < .001. There was no interaction between Solution and Access, *F* < 1. Together, these effects indicate that the GSB and SB groups maintained consistently higher intakes than the GSU and SU groups (i.e. a maintained binge effect), and that despite higher intakes in the GSU and GSB groups on average relative to the SU and SB groups, the binge effects for the saccharin and the mixed glucose and saccharin solution were similar. There was a quadratic trend in intakes across days, *F*(1, 36) = 4.56, *p* = .04. Analyses also revealed a Day by Access linear, *F*(1, 36) = 6.91, *p* = .01, and quadratic interaction, *F*(1, 36) = 4.19, *p* = .048. Figure 5 suggests that the linear interaction can be accounted by increasing intakes in the GSU and SU groups across Phase 2 in contrast to steady intakes in the GSB and SB groups.

#### 4.2.2. Behavioral data

Overall, there were no differences in measures of withdrawal and craving between the Binge and Unrestricted conditions for either of the target solutions.

Almond preferences are shown in Figure 6A. A 2 × 2 × (3) Solution × Access × Test mixed ANOVA failed to detect any main effects of Solution, Access, Test or interaction between these factors (all *p*s > .05).

**Figure 6.**
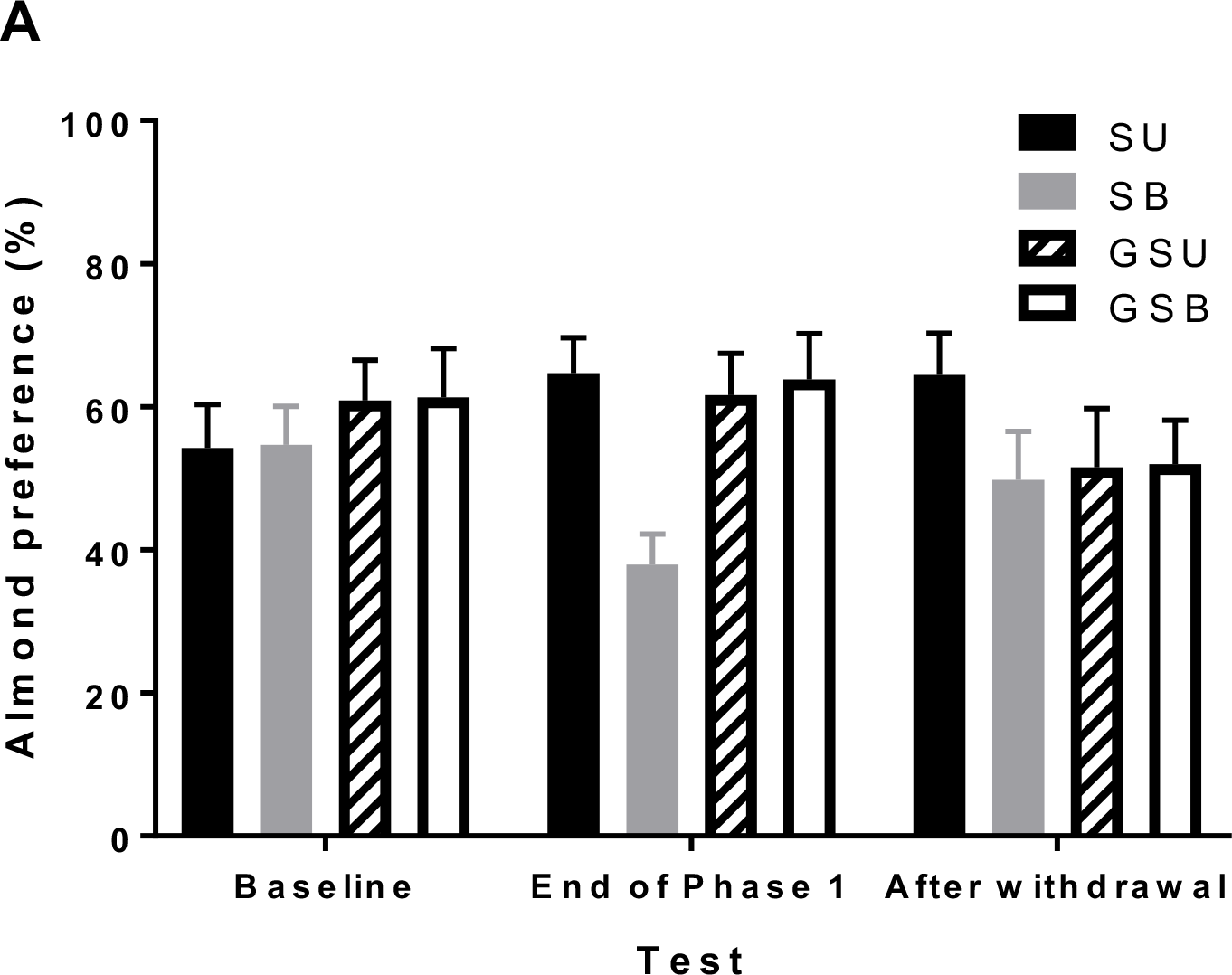
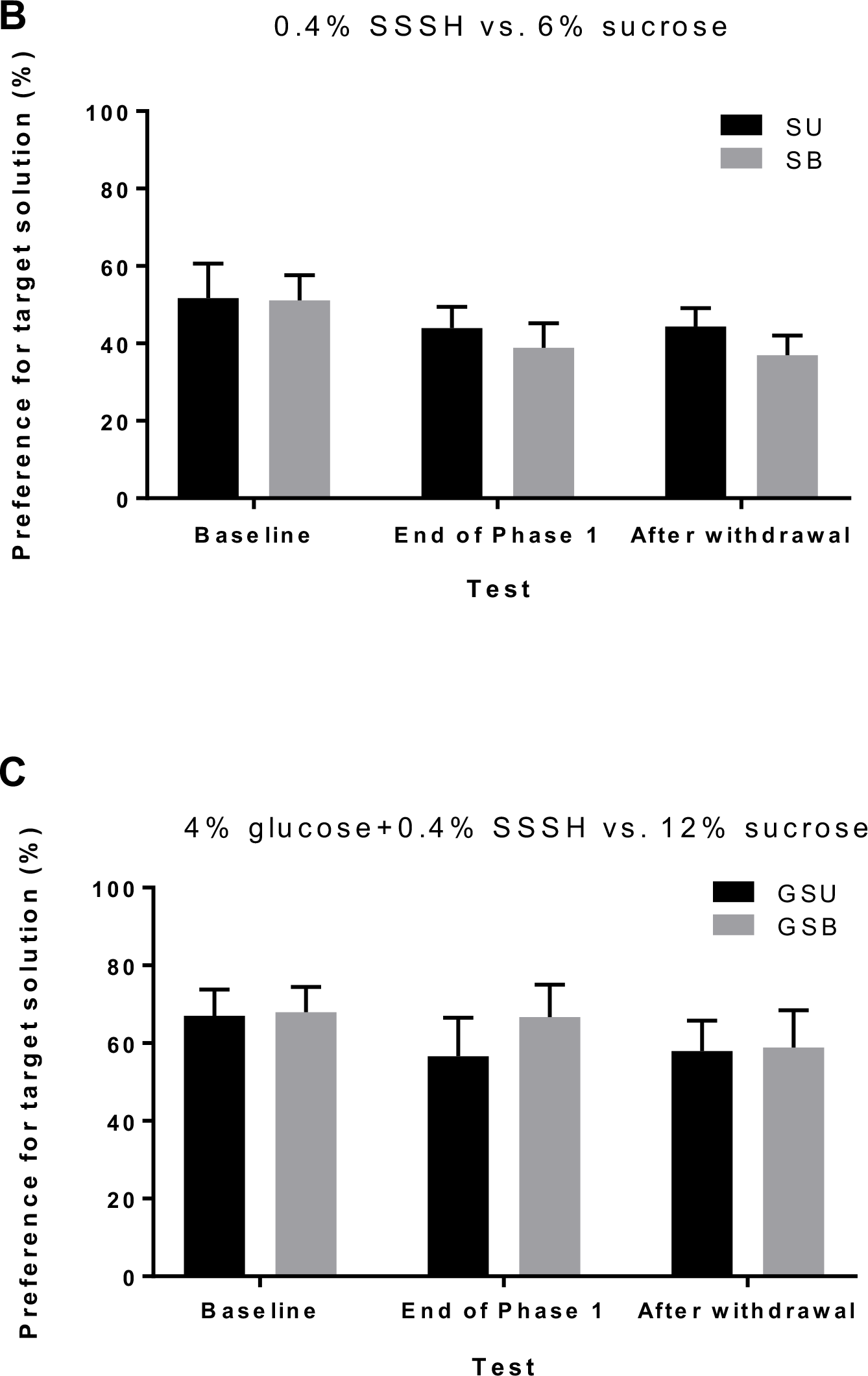
Behavioral data for Experiment 3. A) Mean (± SEM) preference for an almond-flavored solution (1% almond + base) over a flavorless solution (base only) in SU, SB, GSU, and GSB groups. No group differences were found (*p*s > .05) B) Mean (± SEM) preference for 0.4% saccharin (target solution for SU and SB groups) over 6% sucrose solution in a two-bottle choice test conducted at baseline, end of phase 1, and after withdrawal. Preference for saccharin decreased across tests similarly between the SB and SU groups (*p* = .03). C) Mean (± SEM) preference for 4% glucose + 0.4% saccharin (target solution for GSU and GSB groups) over 12% sucrose solution in a two-bottle choice test conducted at baseline, end of phase 1, and after withdrawal. Preferences for the glucose + saccharin solution remained consistent across tests in the GSU and GSB groups (*p*s > .10).

As different concentrations of sucrose were used as the comparison solution for the GSU and GSB groups (12% sucrose) and the SB and SU groups (6% sucrose), 2 × (3) Access × Test mixed ANOVAs were conducted separately for each solution. For the SU and SB groups, there was a main effect of Test, *F*(2, 36) = 4.08, *p* = .03, but no Access effect nor Access by Test interaction (*F*s < 1), indicating that there were no differences in the decrease in preference for saccharin over sucrose across tests between the SU and SB groups (Figure 6B). For the GSU and GSB groups, no main effects nor interactions were found (*p*s > .10), indicating that preference for the glucose + saccharin solution relative to sucrose remained consistent across tests (Figure 6C).

A 2 × 2 between-subjects ANOVA on percentage of open-arm time based on initial scores failed to detect any main effects of Solution and Access, or interaction effect, *F*s < 1. The average percentage of open-arm time was 24.1% in the SU group, 27.3% in the SB group, 29.0% in the GSU group, and 30.4% in the GSB group. These percentages are higher than those obtained in the previous experiment and suggest that the present rats were not anxious. Inter-rater reliability analyses were not run because no group differences were found.

### 4.3. Discussion

The current experiment demonstrated that the persistent binge effect found in Experiment 1 could also been found when using a sweet, but non-caloric saccharin solution and a highly hedonic, mixed glucose and saccharin solution. Both groups receiving every-fourth-day (Binge) access to either saccharin (SB) or glucose and saccharin solution (GSB) escalated their intakes across Phase 1, such that they drank significantly greater amounts in the same 24-h period than the respective groups receiving daily (Unrestricted) access to the same solutions. Most importantly, the differences in intake between rats in the Binge and those in the Unrestricted conditions were maintained across 24 days of Phase 2, despite both groups being switched to identical alternate-day access conditions. Further, although the absolute intakes from the groups given the mixed glucose and saccharin solution were greater than those given saccharin solution alone, the magnitude of the binge effect was similar for the two solutions.

As in the previous experiments the behavioral measures failed to detect any evidence that the binge treatment produced either craving or withdrawal. Despite finding persistent binge-like consumption of saccharin or a mixed glucose and saccharin solution in the SB and GSB groups respectively, almond preference was similar across these groups and their unrestricted counterparts (SU, GSU). This suggests that bingeing on a solution does not increase liking for a flavor paired with that solution. Likewise, when compared to an equally attractive sucrose solution, GSB and SB groups did not prefer their binged target solutions more than the GSU and SU groups. EPM data also showed similar levels of anxiety-like behavior between Binge and Unrestricted rats.

## 5. General Discussion

The current study had two main aims. One was to establish whether the persistence of binge-like consumption induced by the adapted Eikelboom protocol would generalize to similarly attractive solutions. The second was to test whether persistent bingeing would be accompanied by addiction-like behaviors. As discussed in more detail below, the first aim was achieved but no evidence was obtained to indicate that the 1-in-4-days binge treatment produced addiction to any of the solutions used in the three experiments.

Experiment 1 established that 1-in-4-days access to 4% sucrose solution produced an escalation in 24-h intake across exposures and that these elevated intakes persisted when switched to alternate-day access, even when the duration of Phase 1 at 28 days was shorter than the 49 days in Eikelboom and Hewitt (2016) . Experiment 2 extended the adapted Eikelboom protocol by replacing sucrose with two target solutions: 4% maltodextrin and 0.4% saccharin. While 1-in-4-days access to maltodextrin increased intakes, this effect did not persist. Unfortunately, Experiment 2 could not assess whether a persistent bingeing effect could be produced using saccharin. Consequently, Experiment 3 compared 0.4% saccharin solution with a highly palatable mixed 4% glucose and 0.4% saccharin solution. Using these solutions, Experiment 3 found the same persistent bingeing effect as that found for 4% sucrose in Experiment 1, thus satisfying our first aim. As for our second aim, we failed to find ‘withdrawal’ or ‘craving’ in rats engaging in persistent binge-like consumption in all three experiments.

Sweetness appears to be a driving factor in persistent binge-like consumption under the Eikelboom protocol. Our main findings of persistent binge-like consumption of sucrose, saccharin and a mixed glucose-saccharin solution, yet not of maltodextrin, demonstrate the generalizability of the adapted Eikelboom protocol to sweet solutions. In Phase 1 of Experiment 2, 1-in-4-days access to maltodextrin solution in the Maltodextrin Binge (MB) group increased 24-h intakes to levels higher than that of the Maltodextrin Unrestricted (MU) group given continuous access. This finding is consistent with existing studies demonstrating that intermittency increases intakes (Corwin & Babbs, 2012), and our current findings from Experiment 1 and 3. However, when the MB and MU groups were switched to alternate-day access in Phase 2 of Experiment 2, the MU group rapidly increased their intakes to match those of the MB group. As 4% maltodextrin has a similar caloric value to 4% sucrose used in Experiment 1, the collapse of the bingeing effect when maltodextrin was used suggests that caloric value is not critical to the persistent bingeing effect. Supporting this idea, the current study found persistent bingeing using non-caloric saccharin in Experiment 3.

One explanation that Eikelboom and Hewitt (2016) offer for findings of persistent binge-like consumption is that learning about the value of sucrose is different with intermittent access. Mice with a history of daily intermittent access to sucrose or saccharin when food-deprived were later found to exhibit binge-like consumption even after a systematic administration of glucose and chow consumption (Yasoshima & Shimura, 2015). These researchers concluded that intermittent access enhances the hedonic value of a solution rather than induce any homeostatic or metabolic changes (Yasoshima & Shimura, 2015). In the current study, some support for intermittent access increasing hedonic value was found in Experiment 1 using the target solution preference tests; the Binge group had elevated sucrose preferences (relative to maltodextrin) compared to the Unrestricted group at the end of Phase 1. However, no group differences in sucrose preferences were evident at the end of Phase 2 despite the persistent binge effect. On the other hand, Experiment 3 failed to find any group differences in target solution preference (relative to sucrose) at the end of Phase 1 or Phase 2. This inconsistency in finding group differences between experiments even when a persistent binge effect was established suggests that the target solution preference tests may have been insensitive to hedonic changes. However, no direct measure of the hedonic value of the solutions was employed in the present experiments.

The failure to obtain any evidence that the Binge rats became addicted to any of the solutions used in these experiments from the two remaining behavioral measures seems unlikely to be attributed to the inadequacy of the measures used. In Experiment 1, a difference in rate of responding for 4% sucrose was found at the end of Phase 1 between the Chow group and the two sucrose groups, albeit in the unexpected direction whereby the Chow rats responded at a higher rate than the other two groups. There was no suggestion at all of a difference between Binge and Unrestricted groups in lever-press rates at either the end of Phase 1 or the end of Phase 2.

A similar argument applies to the almond preference measure. As seen in Figure 1A, in Experiment 1 almond preferences were higher in the two sucrose groups than in the Chow group at the ends of both Phase 1 and Phase 2 but there was no indication of any difference between the Binge and Unrestricted groups on this measure. As for the data obtained from the almond preference tests in Experiment 3 (see Figure 6A), of the two groups given saccharin the Binge group (SB) displayed lower preferences than the Unrestricted group (SU), while there was no sign of any difference between the two groups given the glucose and saccharin mix (GSU and GSB).

The use of the elevated plus-maze (EPM) to measure possible withdrawal was based on experiments using the Hoebel protocol whereby a higher level of anxiety, as measured on the EPM, was found in some experiments following a period in which intermittent access to a sugar solution was no longer given (Avena et al., 2008). However, it must be noted that in the Hoebel protocol withdrawal-like behaviour was found following a 24-36 h food-deprivation period and/or a naloxone injection (Colantuoni et al., 2002). These conditions were not replicated in the current study. In Experiment 1 the groups did not show any signs of differing levels of anxiety as measured on the EPM. However, the mean percent of time spent on the open arms was low, suggesting that a floor effect - whereby all rats were displaying a high level of anxiety – might have obscured possible group differences. This argument cannot be applied to the EPM results from Experiment 3, where the percentages of time spent on the open arms was higher for all four groups than in Experiment 1 and at a level suggesting a low level of anxiety overall. Thus, as with the craving measures, it seems that the failure to detect a withdrawal effect in the Binge groups was unlikely to be because of insensitivity of the measure employed. Rather, previous reports of addiction-like behaviours accompanying bingeing may not exist under the more controlled and circadian-independent conditions of the current study protocol.

The prediction that addiction-related effects would be produced by the present procedures was partly based on the evidence obtained from what we have referred to as the Hoebel protocol, whereby rats are given 12-h access each day to a sugar solution (Avena et al., 2008). It may be noted that a recent substantial study that used 10% sucrose in this protocol found that the procedure reduced, rather than increased, *wanting* for the sucrose solution. The measure used in these experiments was a conditioned place preference test, which is analogous to the almond preference measure used here (Smail-Crevier, et al., 2018). It is also worth mentioning that in comparison to multiple studies focusing on a single outcome measure using the Hoebel protocol (e.g. Avena, Bocarsly, Rada, Kim, & Hoebel, 2008; Avena & Hoebel, 2003; Avena et al., 2005; Colantuoni et al., 2002), the current study used multiple outcome measures of addiction-like behaviours in the same experiment to fully assess whether access-induced bingeing behaviour can be appropriately considered ‘addictive’.

In conclusion, these experiments indicate that the hedonic value of a solution is more important than its caloric value in determining whether 1-in-4-days intermittent access to a solution will produce persistent bingeing. However, they suggest that such persistence is not produced by some kind of addiction to the solution, since our assessments of withdrawal and compulsive-like behaviour toward the putative addictive substance in each case yielded null results.

## Acknowledgements

This study was partly supported by Australian Research Council Discovery grant DP170103927 and partly by the School of Psychology at the University of Sydney. Experiments 1 and 2 were reported by SR in an Honours thesis in Psychology (2017). We are grateful for the technical help provided by Nenad Petkovski and for advice on procedures plus comments on the manuscript by Dr. Michael Kendig.

